# RNA folding kinetics control riboswitch sensitivity in vivo

**DOI:** 10.1101/2024.03.29.587317

**Authors:** David Z. Bushhouse, Jiayu Fu, Julius B. Lucks

## Abstract

Riboswitches are ligand-responsive gene-regulatory RNA elements that perform key roles in maintaining cellular homeostasis. Understanding how riboswitch sensitivity is controlled is critical to understanding how highly conserved aptamer domains are deployed in a variety of contexts with different sensitivity demands. Here we uncover new roles by which RNA folding dynamics control riboswitch sensitivity in cells. By investigating the *Clostridium beijerinckii pfl* ZTP riboswitch, we identify multiple mechanistic routes of altering expression platform sequence and structure to slow RNA folding, all of which enhance riboswitch sensitivity. Applying these methods to riboswitches with diverse aptamer architectures that regulate transcription and translation with ON and OFF logic demonstrates the generality of our findings, indicating that any riboswitch that operates in a kinetic regime can be sensitized by slowing expression platform folding. Comparison of the most sensitized versions of these switches to equilibrium aptamer:ligand dissociation constants suggests a limit to the sensitivities achievable by kinetic RNA switches. Our results add to the growing suite of knowledge and approaches that can be used to rationally program cotranscriptional RNA folding for biotechnology applications, and suggest general RNA folding principles for understanding dynamic RNA systems in other areas of biology.

## INTRODUCTION

Riboswitches are RNA sensors that regulate gene expression based on the cellular concentration of their cognate ligands (1). Riboswitches comprise two functional domains, a ligand-binging aptamer domain (AD) and a gene-regulatory expression platform (EP), which interface to enable ligand-dependent cis-regulation of downstream genes. Across bacteria, riboswitches are integrated into diverse gene-regulatory circuits that maintain cellular homeostasis by sensing a wide variety of metabolites, ions, enzyme cofactors, amino acids, uncharged tRNAs, and other signaling molecules (2, 3). Because they control central housekeeping functions within pathogenic organisms, there is growing exploration of riboswitches as antibiotic targets (4–7). Riboswitches have also served as excellent model systems to study RNA structure/function relationships (8–14), and increased understanding of riboswitch mechanisms has enabled them to be engineered in biotechnology applications (15–17), most notably as biosensors for environmental (18, 19) and human health biomarkers (20, 21).

A central goal of riboswitch biology is to understand the molecular mechanisms used by these ubiquitous RNA sensors to make gene-regulatory decisions. While much attention has been focused on understanding how properties of the AD, such as the K_D_ of ligand binding, govern riboswitch function (22), recent studies have found that ligand-binding interactions alone are poor predictors of riboswitch regulatory potential (23–25). These findings reveal shortcomings in our current understanding of the biochemical principles underlying riboswitch functional properties, namely sensitivity (EC_50_) and dynamic range.

Broadly, it has been well known that transcriptional dynamics, including transcription speed and RNAP pausing, can affect riboswitch function by altering the RNA cotranscriptional folding landscape (26–30). However, recent work is showing that the RNA folding kinetics of the EP itself can influence function as well. Recently, we showed that introducing kinetic barriers to EP folding via internal strand displacement resulted in quantitative increases in the dynamic range of the *Clostridium beijerinckii pfl* (*Cbe pfl*) ZTP riboswitch, demonstrating that EP RNA folding kinetics can also control riboswitch function (31). However, it is an open question whether EP folding can control riboswitch sensitivity, or if sensitivity is solely governed by the ligand-binding AD domain.

Here, we investigate this question through the detailed study of several classes of transcriptional riboswitches. Transcriptional riboswitches regulate transcription elongation, which requires these riboswitches to fold into a ligand-binding competent AD structure, sample the environment for ligand, and undergo ligand-dependent global structural rearrangements to produce a genetic decision, all during active transcription. Transcriptional riboswitches have been demonstrated to operate in a kinetic regime, meaning that they feature a ligand-binding time window that kinetically constrains the ligand:AD binding interaction before the gene-regulatory decision is made (22, 27, 28, 32). This means that all transcriptional riboswitches must execute their genetic decision within the ms-s timeframe before RNAP escapes the termination site (33). Given this feature, we hypothesized that increasing the ligand-binding time window through altering EP folding kinetics could serve to enhance the sensitivity of kinetically operated riboswitches.

We first investigated this hypothesis through an in vivo functional mutagenesis approach applied to the *Cbe pfl* ZTP riboswitch. *Cbe pfl* is an ideal model system, as its switching mechanism features a well-characterized 7-10 nt ligand-binding time window extending from the initial formation of the ligand binding-competent AD to the nucleation of the EP via the closing of an RNA hairpin loop that ultimately results in the formation of an intrinsic terminator hairpin (Fig. S1A) (10, 34, 35). We found that altering the length and sequence of the *Cbe pfl* EP terminator loop finely controls riboswitch sensitivity. Extending this single-stranded region enhances EC_50_, as does the presence of A-A stacking interactions near the 3’ end of this domain.

Unexpectedly, we discovered a functional tradeoff between sensitivity and dynamic range, which we show by systematic mutational analysis to be mediated by A-A stacking within the EP loop region. Functionally characterizing 3,289 different EP loop sequences using a high-throughput FACS-seq approach showed that this tradeoff is inherent to the *Cbe pfl* EP architecture. Applying a synthetic approach, we designed a new remote toehold EP architecture that escapes this functional tradeoff, enabling simultaneous enhancement of sensitivity and dynamic range.

The discovery of the remote toehold mechanism prompted us to investigate whether it is present in nature. To explore this, we bioinformatically investigated the diversity of natural ZTP riboswitch EPs, which revealed a range of EP architectures. Functional characterization of a sample of these architectures confirmed that diverse structural approaches are used in nature to tune sensitivity, all involving changes to the EP that are predicted to alter EP folding pathway kinetics.

We next sought to generalize our findings by investigating other riboswitches. We began with a version of the ZTP riboswitch from *Pectobacterium carotovorum* that regulates translation. We found that introducing kinetic barriers to EP nucleation enhanced sensitivity in this riboswitch as well, indicating that translational ZTP riboswitches can also operate in a kinetic, cotranscriptional regime. With rational changes we then mutated this riboswitch to control transcription, showed that it can also be sensitized through similar EP changes, and found that transcriptional and translational variants approach the same limit of sensitivity independent of EP architecture.

Finally, we applied principles learned from ZTP riboswitches to rationally design EP mutations for a range of transcriptional riboswitches from other classes, including the *pbuE* and *yxjA* purine riboswitches and the *crcB* fluoride riboswitch, and showed that these mutations all resulted in sensitization with trends that match those found in the ZTP system. A comparison between the most sensitive versions of these classes and their equilibrium AD:ligand K_D_ values show that there is a limit to the sensitivity that can be achieved with kinetically driven switches, suggesting that in vivo cotranscriptionally folded aptamer domains access structural states that have weaker ligand affinities than those characterized with refolded RNA in vitro.

Taken together, these results suggest a general principle by which EP folding kinetics can control riboswitch sensitivity for diverse riboswitches that operate in a kinetic regime. This finding may help explain the diversity of natural riboswitch EPs, give insights into riboswitches as antibiotic targets, and provide new routes to rationally engineering riboswitches for a range of applications. Given that diverse non-coding RNAs have also been shown to utilize kinetically driven folding pathways similar to riboswitches (36), these results add to our generalized understanding of how dynamic RNA folding can impact RNA function (37).

## METHODS

### Cloning and plasmid construction

Riboswitch reporter plasmids were prepared as described previously (31). Briefly, plasmids were constructed on a p15A plasmid backbone with a reporter cassette comprising a J23119 σ70 consensus promoter, riboswitch variant, ribosome binding site, and superfolder green fluorescent protein (sfGFP) coding sequence. Riboswitch mutants were generated by inverse polymerase chain reaction (iPCR) followed by blunt end ligation. Sequences were confirmed by Sanger sequencing (Quintara Biosciences), and plasmid stocks were screened for oligomerization by agarose gel electrophoresis. Approximately 20% of clones generated by the iPCR method were visibly oligomerized and discarded. Correct constructs were re-transformed into 10-beta chemically competent cells (NEB), from which clones were used to prepare glycerol stocks (25% glycerol, stored −70°C) and minipreps for data collection. Sequences of all reporter plasmids can be found in Supplementary Data File 1 along with Addgene accession numbers for select constructs.

### In vivo bulk fluorescence reporter assay

Riboswitch functional assays were performed as described previously (31). Briefly, *E. coli* BW25113 (Keio parent) was transformed with reporter plasmids, and colonies were used to inoculate 300 μL overnight cultures in LB media. 4 μL of overnight cultures were then used to inoculate 200 μL subcultures in M9 enriched media (1x M9 salts, 1 mM thiamine hydrochloride, 0.4% glycerol, 0.2% casamino acids, 2 mM MgSO_4_, 0.1 mM CaCl_2_, 34 μg/ml chloramphenicol) containing the indicated amount of ligand. For ZTP riboswitch assays, 5-aminoimidazole-4-carboxyamide-ribonucleoside (Z) (Millipore Sigma) in DMSO was used; for purine riboswitch assays 2-aminopurine (2AP) (Millipore Sigma) in DMSO was used. For fluoride riboswitch assays, reporter plasmids were transformed into *E. coli* JW0619 (Δ*crcB* Keio), and subcultures were treated with NaF (Millipore Sigma) in water. Subcultures were shaken at 1000 rpm and 37°C in a benchtop incubator for 6 hr, after which sfGFP fluorescence and OD_600_ were measured on a Biotek Synergy H1 microplate reader. All in vivo reporter assays were performed with three experimental replicates, each performed in triplicate with three separate colonies (biological replicates) for a total of nine data points (n = 9) per reporter construct. To obtain sensitivity and dynamic range information, each construct was treated with a range of eight ligand concentrations. Therefore, each reported EC_50_ and fold change value is the result of 72 independent measurements comprising three experimental replicates, triplicate biological replicates, and eight different ligand conditions.

Fluorescence data was processed by subtracting average media blank measurements from both OD_600_ and fluorescence measurements, scaling fluorescence values from arbitrary units to micromolar equivalent fluorescein (μM fluorescein, MEF), and dividing fluorescence values by OD_600_ values for each sample containing well. The fluorescein scaling factor was determined by performing a calibration procedure on the microplate reader used to collect all fluorescence data. To convert fluorescence values, measured arbitrary units were divided by the fitted slope of this calibration curve without subtracting the fitted intercept. More information about fluorescence calibration is available in Supplementary Data File 1. Dose response data from the 72 independent measurements was analyzed by performing least squares fitting (code available at https://github.com/LucksLab/Bushhouse_Riboswitch_Sensitivity_2024) to the collection of data points using the Hill equation:

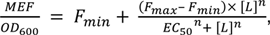

where [L] is the concentration of ligand supplemented to the media, the fit parameters F_min_ and F_max_ account for baseline expression (leak) and the theoretical maximum gene expression under saturating ligand concentrations (ON-state), and the fit parameter EC_50_ is the concentration of supplemented ligand required to induce half-maximum fluorescence. The Hill coefficient n was heuristically set based on the riboswitch class: n = 1.35 for ZTP riboswitches, n = 1 for *pbuE* riboswitches, n = −1 for OFF ZTP riboswitches, n = 2.2 for fluoride riboswitches, and n = −1.45 for *yxjA* riboswitches. We performed this procedure to ensure all of the error of the fits was contained in the EC_50_ and fold change values. See Fig. S2-3 for more detail about the fitting procedure. EC_50_ values were extracted from fitting to the 72 data points. Fold change was calculated by dividing the fitted F_max_ by F_min_, with standard error propagation:

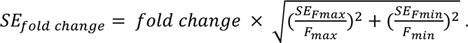

All fluorescence data and the resulting fits are available in Supplementary Data File 1. Where reported, EC_50_, F_min_, F_max_, and fold change are from fits to these 72 independent measurements with errors reported as standard error of the fits or propagated errors as described. All plots were generated using DataGraph 5.3.

### Bioinformatic analysis of ZTP riboswitch EPs

1755 putative ZTP AD sequences from Rfam entry RF01750 were used to query the relevant genomes in GenBank for the downstream 200 bp. These putative riboswitch-containing loci were tested for the presence of a polyU tract (6 Ts in an 8 nt sliding window) and an Rfam consensus P3 stem. Secondary structure prediction (ViennaRNA 2.6.3) was performed on the subdomains extending from P3 to the polyU tract to ensure the formation of a single terminator stem-loop structure. The EP was examined for the presence of a region complementary to the 3’ side of the Rfam consensus P3 stem, which was marked as a putative invader. Terminator-containing EPs were manually classified using NUPACK secondary structure prediction to validate invader predictions and annotate EP architecture (loop-only, direct toehold, remote toehold). All bioinformatic code was written using ChatGPT 3.5 (OpenAI), and is available at https://github.com/LucksLab/Bushhouse_Riboswitch_Sensitivity_2024.

### Single round in vitro transcription functional assay

Transcription termination assays were performed as described previously (31). Briefly, 30 second single round IVT reactions were performed with 400 μM NTPs in the presence or absence of the indicated ligand concentration. Reaction products were isolated by phenol-chloroform extraction and DNase digestion, run on denaturing polyacrylamide gels, stained, and imaged for quantification. IVT assays were performed in duplicate.

### FACS-seq high throughput assay

A FACS-seq high throughput assay was developed based on existing methods (38–40). A library of *Cbe pfl* reporter plasmids with random terminator loop sequences was generated by performing iPCR cloning with a hand-mixed random primer (IDT). Ligated PCR products were electroporated into *E. coli* BW25113 and plated on several LB agar plates overnight at 37°C. The following morning, plates were transferred to the benchtop for approximately 8 hr, after which colonies were combined by repeatedly pipetting 10 mL of LB across the surface of each plate and resuspending colonies with sterile spreader arms. The combined resuspended library was vortexed briefly, passed through a 100 μM filter (CELLTREAT) to remove cell aggregates, and used to inoculate a 5 mL LB overnight culture shaken at 37C. The following morning, 100 μL of overnight culture was used to inoculate 5 mL subcultures in M9 enriched media containing the appropriate amount of Z. Subcultures were shaken at 37C for 6 hr, after which they were passed through a 100 μM filter (CELLTREAT), diluted 50x in 1x PBS, and placed on ice for 2 hr. Library subcultures were preliminarily characterized using a BD Accuri ™ C6 Plus flow cytometer to examine library fluorescence as a function of supplied Z (Fig. S9B).

A sample of each subculture was sorted into 4 bins based on FITC fluorescence intensity (Fig. S9B). Sorting was performed using a BD FACSMelody™ Cell Sorter at the Robert H. Lurie Comprehensive Cancer Center Flow Cytometry Core Facility at Northwestern University. Cell sort counts for each bin of each subculture were recorded and used in the data analysis procedure below (Supplementary Data File 1). After sorting, cells were placed on ice for 2 hr, after which the entire volume of sorted cells for each bin, as well as 1 mL of unsorted diluted subculture for each ligand condition, were used to inoculate separate 5 mL LB cultures, which were shaken overnight at 37°C and miniprepped.

Sequencing libraries for each sorted bin were generated by two-step PCR using a Phusion® DNA Polymerase (NEB) kit. In the first round, for each sample, a 100 μL PCR was assembled using 1 μg of input plasmid template and 400 nM of both Round 1 primers (DZB.C08, DZB.C09) which were designed to install 5 nt of random sequence on both sides of the amplified region to increase sequence diversity during initial sequencing cycles. After 3 rounds of PCR, reactions were placed briefly on ice, and 0.5 μL of ExoI (NEB) was mixed into each reaction. Reactions were incubated at 37°C for 30 min to digest Round 1 primers, and 80°C for 20 min to denature ExoI. Following ExoI deactivation, Round 2 primers (Illumina TruSeq Adapters) were added to each reaction to 400 nM, and PCR was resumed for 10 additional cycles. A unique i7 index was used for each sorted bin (Supplementary Data File 1). PCR products were purified from template plasmid and primers by two-step bead purification using SPRI beads (Cytiva) and quantified using Qubit 3.0 High Sensitivity dsDNA kit (Invitrogen). Indexed libraries across ligand and bin conditions were pooled in equimolar ratios and submitted to the NUSeq Core Facility for sequencing on a full flowcell of a NextSeq 500 (Illumina) instrument for paired-end 75 x 75 cycles, with an expected 58 bp of overlap between read 1 and read 2. The entire FACS-seq experiment from library construction to sequencing was performed in duplicate. Sequences of oligos used for sequencing library preparation can be found in Supplementary Data File 1.

For each ligand condition, the FACS-seq protocol generated 4 pairs of paired-end sequencing files, each containing the reads from one of the four sorted bins. Sequencing reads were analyzed to generate mean fluorescence estimates for each loop sequence in each ligand condition (40). Dose response curves were then fit across ligand conditions to determine functional properties (EC50, fold-change) for each loop sequence (Fig. S9). Data was processed as follows:

Paired-end sequence files for each index were first filtered to remove reads with Qscore < 30 and reads in which the 58 bp of overlapping sequence in the forward and reverse reads did not match. Reads that failed to match the riboswitch template sequence or contained ambiguous base calls (‘N’) were also discarded.

The loop sequences from all remaining reads were extracted and counted based on the index, bin and ligand conditions, resulting in a table of read counts for each loop sequence, *i*, in each bin *j*, in each ligand condition,*k*, *read count*_*i*,*j*,*k*_. The FACS sorting process results in different numbers of sorted cells in each bin and ligand condition. We therefore normalized read counts according to:

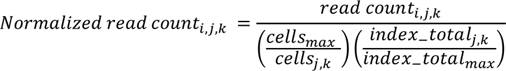

where *index_total_j_* and *cells_j_* are the number of reads and the number of cells sorted for a given bin *j* and ligand condition *k*, respectively, and *index_total_max_* and *cells_max_* are the maximum number of total reads and the number of cells sorted across bins and ligand conditions, respectively. The factor 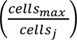 reduces the signal from bins that have a low number of sorted cells which would normally be over-represented, while the factor 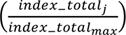 boosts the signal from bins that had lower numbers of reads sampled during sequencing which would normally be under-represented.

For each loop sequence, mean fluorescence at each ligand condition (*F*_*i*,*k*_) was estimated by calculating the weighted average of the normalized read counts across the four sorted bins:

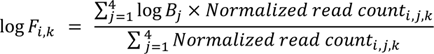

where *B_j_*denotes the geometric mean fluorescence value of the bin *j*, calculated using the upper, *U*_*j*_, and lower, *L*_*j*_, bounds of the gate applied during cell sorting (Supplementary Data File 1):

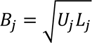

Mean fluorescence values were calculated for each loop sequence in each ligand condition, converted out of log space, and values of both experimental replicates were pooled before fitting dose-response curves using the Hill equation as described above. Data quality criteria were applied to remove poor fits and loop sequences with sparse data. Loop sequences with reads in fewer than 3 bins for any ligand concentration in both replicates were excluded, as were loop sequences where the ratio of the standard error of the fit parameters to the parameters themselves 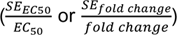 exceeded 1, or if the extracted EC_50_ value exceeded 500 μM. Of the 4912 loop sequences with reads in ≥ 3 bins at each ligand concentration in at least one replicate, 3289 passed the data quality criteria and were used for subsequent analysis. Density plots were generated by performing a gaussian kernel density estimation on the fold change and EC_50_ measurements of the 3289 characterized loops. All FACS-seq data analysis code was written using ChatGPT 3.5 (OpenAI) and is available at https://github.com/LucksLab/Bushhouse_Riboswitch_Sensitivity_2024.

## RESULTS & CONCLUSIONS

### Cbe pfl terminator loop length and sequence tune riboswitch sensitivity

The *Cbe pfl* ZTP riboswitch AD ligand binding pocket is formed by a pseudoknot between the P3 stemloop an upstream helix-junction-helix motif (41). This ligand-binding competent structure is mutually exclusive with the formation of the EP, which consists of a long intrinsic terminator involving all of the nucleotides from P3 (35) (Fig. S1B-C). The folding pathways of three ZTP riboswitches have been biophysically characterized (10, 34, 35). Together, these studies indicate that during transcription, the ligand-binding window opens shortly after the 3’ end of the P3 stemloop emerges from the RNAP exit channel, remains open during the transcription of the downstream ∼7-10 nt (including the EP loop sequence), and closes when the first 2-3 nts of the EP invader domain nucleate the strand displacement of P3, after which the AD loses the ability to bind ligand (34) (Fig. S1A). Therefore, the time constraint posed by this short ligand-binding window should be determined by the amount of time required to transcribe the EP loop sequence between P3 and the invader domain, as well as the rate of the RNA folding step that nucleates EP strand displacement. We hypothesized that both of these factors could be affected by the EP loop length, since longer loops face an increased entropic cost to close (42).

To test this hypothesis, we varied the 7 nt polyA loop of *Cbe pfl* (Fig. 1A) from 3 nt to 16 nt, and altered the sequence from polyA to polyU or polyC. We measured dose-response curves in vivo, and fit them to the Hill equation (see **Methods**), allowing us to characterize sensitivity (EC_50_), and fold change (ratio of the maximum gene expression activation to the minimum) for each variant. We observed that extending homotypic EP loops enhances riboswitch sensitivity (Fig. 1B-C), with polyA loops resulting in lower EC_50_ relative to polyU/C loops of the same length (Fig. 1B). While any loop sequence should slow initial EP nucleation as a function of length by increasing the entropic cost of loop closure, we reasoned that polyA stacking interactions within the loop may slow the rate of loop closure further by introducing an additional energetic barrier (43–45). Unsurprisingly, loop extension also resulted in increases in uninduced expression (leak, F_min_) (Fig. S4A-C), since longer loops decrease hairpin stability (46), thus affecting termination efficiency (Fig. S4D).

**Figure 1:**
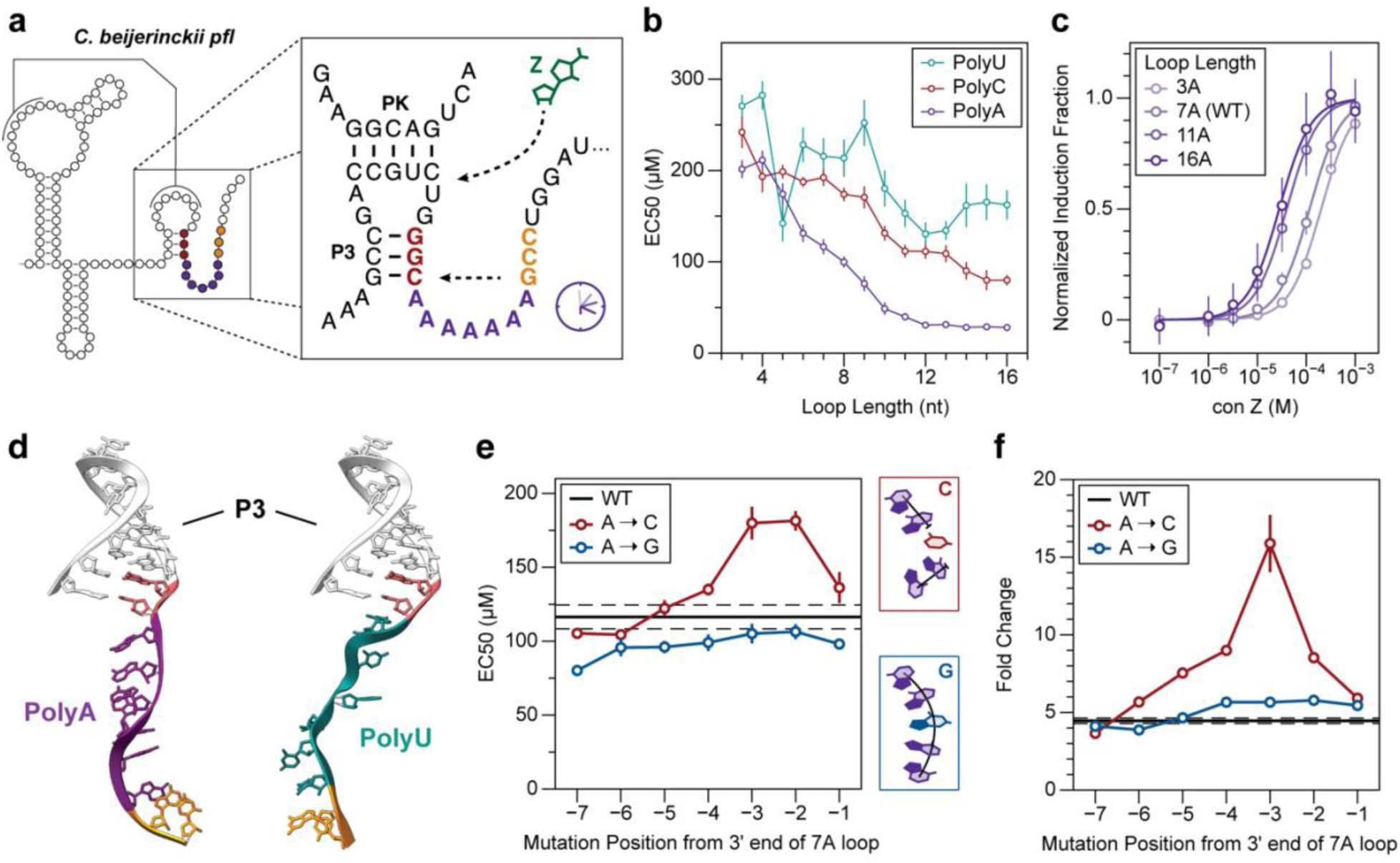
*Cbe pfl* terminator loop length and sequence tune riboswitch sensitivity. (a) Secondary structure schematic of the *Cbe pfl* riboswitch immediately preceding EP nucleation. The polyA terminator loop (purple) closes to allow the invader domain (gold) to initiate strand displacement of the P3 stemloop (red). Ligand binding kinetically competes with this process to determine the riboswitch genetic decision. (b) EC_50_ values for riboswitch variants with homopolymeric loops of varying lengths. (c) Normalized dose response curves of riboswitch variants with varying polyA loop lengths. (d) Vfold3D models of P3 with un-nucleated 7nt polyA or polyU terminator loops. PolyA (purple, left) exhibits regular stacking that separates the invader nucleotides (gold) from the substrate nucleotides (red) in the P3 stem, whereas polyU (teal) exhibits no regular order. (e) EC_50_ values for riboswitch variants with length 7 polyA loops in which one residue has been mutated to either C (red) or G (blue). Nucleotide coordinates are relative to the 3’ end of the loop. Solid horizontal line indicates EC_50_ value for WT polyA, with dashed horizontal lines indicating the standard error. Cartoon insets illustrate how A→C mutations (top) will disrupt local stacking, while A→G mutations allow local stacking interactions (bottom). (f) Fold change of the mutants in (e). Solid horizontal line indicates fold change of WT polyA, with dashed horizontal lines indicating the standard error. Data in panel (c) indicates normalized dose response curves over n = 9 replicates, with error bars indicating standard deviation. Dose response curves were normalized by dividing all data points for each mutant by (F_max_ – F_min_) of the fitted curve, with standard deviation calculated from the 9 normalized replicate values for each concentration. Data in panels (b), (e), and (f) are determined as described in **Methods**.

To visualize how base stacking in the polyA loop might slow strand displacement nucleation, we performed 3D structure prediction of polyA and polyU un-nucleated loops using Vfold3D (47). We observed that the polyA loop (purple) stacks neatly, pointing the invading nucleotides away from the P3 stem, while the polyU loop (teal) is modeled to be unstructured (Fig. 1D). We therefore hypothesized that disrupting stacking interactions within the polyA loop would decrease sensitivity by removing the energetic barrier to loop closure. To test this hypothesis, we performed a mutational scan of the 7 nt polyA loop in which each residue was mutated to C, which should destabilize neighboring A-A stacking interactions (48, 49). We observed that A→C mutations near the 3’ end of the loop region resulted in desensitization (Fig. 1E). Performing the same mutational scan with A→G mutations, which should maintain neighboring stacking interactions, showed that these mutations did not affect sensitivity (Fig. 1E), supporting our hypothesis that stacking interactions within the EP loop enhance riboswitch sensitivity by slowing loop closure kinetics. We further tested these trends by chimerically replacing the 7A loop of *Cbe pfl* with sequence variants of the 12 nt A-rich domain from the *B. subtilis queC* preQ1 riboswitch (*Bsu queC*) found by NMR to have A-A stacking interactions that rigidify the domain (48), and observed similar trends (Fig. S5). These results support the conclusion that polyA stacking within the loop region, especially near the 3’ end, extends the ligand-binding window by slowing strand displacement nucleation.

Surprisingly, the A→C disruptions to stacking in the 7A polyA loop that resulted in desensitization also resulted in marked increases in dynamic range, while A→G mutations had neither effect (Fig. 1F). The same relationship was observed in the *Bsu queC* mutants (Fig. S5B-C), suggesting that the effects of disruptions to A-A stacking on sensitivity and fold-change are somehow coupled in an apparent tradeoff relationship.

Taken together, these results support our hypothesis that EP loop length and sequence composition controls riboswitch function, likely through posing kinetic barriers (entropic or energetic) to loop closure that extend the ligand-binding window. However, disrupting EP loop stacking has the apparently coupled effect of decreasing sensitivity and increasing dynamic range, which we sought to investigate further.

### PolyA loop stacking mediates tradeoff between sensitivity and dynamic range

To interrogate the mechanistic basis by which disruptions to A-A stacking interactions within the *Cbe pfl* EP terminator loop result in apparently coupled increases in EC_50_ and fold change, we performed A→C mutational scans on a variety of polyA loop lengths, and observed similar phenotypic patterns regardless of overall loop length (Fig. 2A-C). Consistently, A→C mutations near the 3’ end of the loop had the greatest effects in simultaneously decreasing sensitivity and increasing fold change (Fig. 2B-C). Enhancements in fold change were caused by dramatic position-dependent increases in ON-state gene expression for these mutants (Fig. S6A-F). Previous work in the ZTP system showed that disfavoring the EP strand displacement process by disrupting the C100:G108 nucleating base pair results in similar dramatic increases in ON-state gene expression (31), suggesting that disrupting A-A stacking interactions within the terminator loop may also disfavor downstream strand displacement.

**Figure 2:**
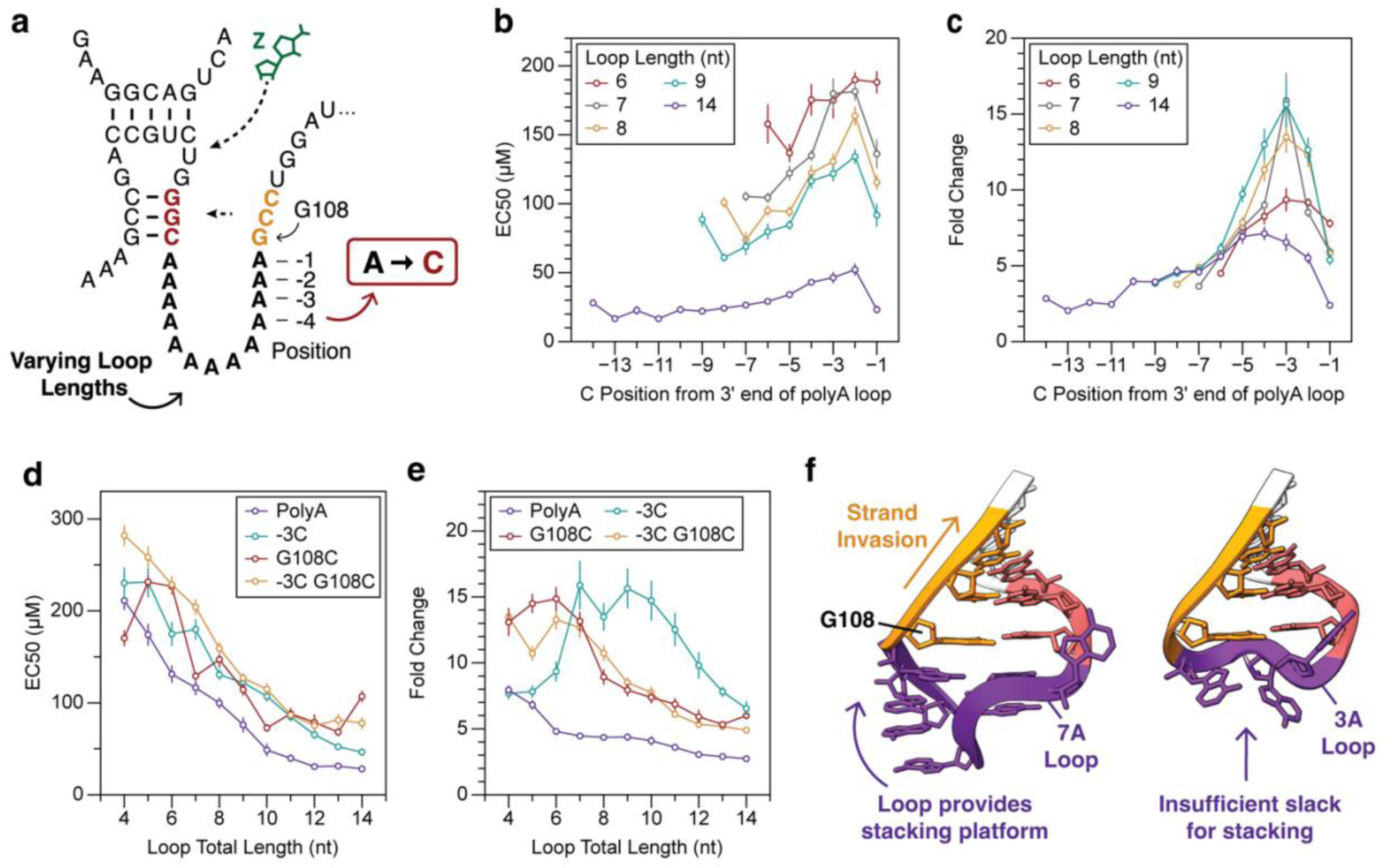
PolyA loop stacking mediates tradeoff between sensitivity and dynamic range. (a) Secondary structure schematic of A→C mutational scans in which riboswitch variants with different length polyA loops were mutated at each position within the loop from A to C to disrupt local A-A stacking. Positional coordinates refer to the distance from the 3’ end of the loop. (b) EC_50_ measurements for mutational scans of different length terminator loops into which single A→C mutations have been made at every position. (c) Fold change of the mutational scans in (b). (d) EC_50_ measurements of riboswitch variants with polyA loops of varying lengths into which either no mutations have been made (polyA), the third to last residue relative to the 3’ end of the loop has been mutated to C (−3C), the initial invading residue G108 has been mutated to C (G108C), or both mutations have been made (−3C G108C). (e) Fold change of the mutational scans in (d). (f) Vfold3D models of the nucleated *Cbe pfl* terminator loop featuring either a length 7 or length 3 polyA loop. The longer loop allows continuous stacking of A residues within the 3’ end of the loop and the G108 invader residue, while the shorter loop does not allow enough slack for stacking to occur in this conformation. Additional loop length models in Fig. S7. Data in (b), (c), (d) and (e) are determined as described in **Methods**.

To test this hypothesis, we extended loop length in the presence of the G108C mismatch mutation (G108C), an A→C mutation at the −3 position relative to the 3’ end (−3C), or both −3C and G108C (−3C G108C). While all of the loop extensions resulted in similar desensitization compared to perfect polyA across loop lengths (Fig. 2D), we observed key differences in the fold change phenotypes between these mutational series (Fig. 2E, S6G-I). While G108C enhances fold change regardless of loop length as expected, −3C mutations do not enhance fold change for loops shorter than 6 nt (Fig. 3E). At the same time, perfect polyA loops shorter than 6 nt show enhanced fold change relative to longer loops, caused by large increases in ON-state gene expression (Fig. 3E, S4A). Taken together, these findings suggest that a minimum loop length of around 6 nts is required for 3’ A-A stacking to suppress fold change.

**Figure 3:**
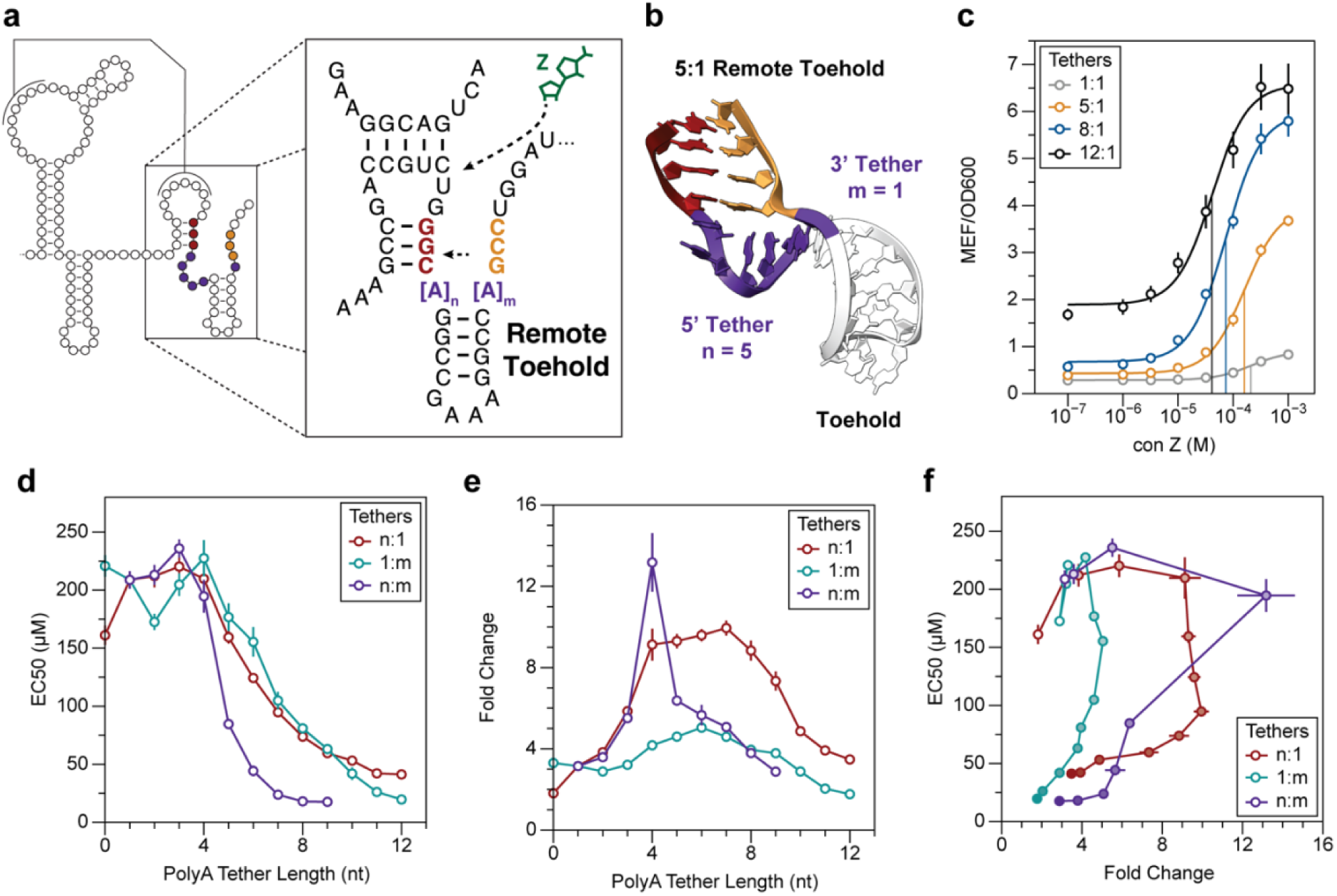
Remote toehold EP architecture uncouples tradeoff between sensitivity and dynamic range. (a) Schematic of the remote toehold EP design, in which the polyA loop of *Cbe pfl* is replaced by a synthetic hairpin connected to the AD and invader by single stranded extendable polyA tethers. (b) Vfold3D model of a nucleated terminator featuring a remote toehold extension with a 5A 5’ tether and a 1A 3’ tether (5:1). (c) Dose responses of select remote toehold variants with varying length 5’ tethers, including the 5:1 design shown in (b). Vertical lines indicate EC_50_ values of the curve with the corresponding color. (d) EC_50_ measurements for remote toehold variants with varying length 5’ and 3’ tethers. n:1 indicates the length of the 5’ tether has been extended, with the length of the 3’ tether fixed at 1A. (e) Fold change measurements for variants in (d). (f) EC_50_ and fold change data from (d) and (e) plotted against each other. Increasing opacity represents longer overall tether length. Data in panel (c) are dose response curves over n = 9 replicates, with error bars indicating standard deviation. Data in panels (d), (e), and (f) are determined as described in **Methods**.

To try to understand why a minimum loop length is required, we performed 3D modeling of the nucleated terminator hairpin with polyA loops of varying lengths using Vfold3D. While this modeling approach is limited in its ability to assess the molecular dynamics of single stranded regions, it revealed a putative stacking platform of A residues within the 3’ side of the terminator loop, which was not present in loops shorter than 6 nt (Fig. 2F, S7). Stacked residues within this 3’ region may align with G108 to provide an energetic backstop to reverse branch migration, analogous to the ‘cooperative toehold’ approach in DNA nanotechnology (50), increasing the overall forward rate of strand displacement and decreasing ON-state gene expression. Shorter loops had insufficient slack to align multiple parallel purine rings below G108 (Fig. 2F, S7), explaining the observation that the fold change of short polyA loop variants is not affected by −3C mutations (Fig. 2E). Importantly, we observe that −3C in the G108C background results in no additional increase in fold change regardless of loop length (Fig. 2E). This apparent epistasis of G108C over −3C is consistent with the hypothesis that a 3’ A-A stacking platform in the terminator loop stabilizes G108 to enhance strand displacement efficiency.

Previous studies have found that toeholds, base paired regions preceding strand displacement initiation sites, are potent enhancers of strand displacement efficiency (51). To test if a toehold would suppress riboswitch fold change in the same manner as 3’ A-A stacking, we performed a mutational series in which a length 13 polyA loop was progressively shortened with A→U mutations, converting loop sequence into toehold base pairs, either in the A:U or U:A orientation (Fig. S8A). Both 5’ and 3’ mutational series resulted in progressive desensitization, as initial loop nucleation should be made much easier by the presence of a toehold region (Fig. S8B). However, we observed that while 5’ A→U mutations that preserve 3’ A-A stacking decreased fold change (Fig. S8C, E), 3’ A→U mutations that break 3’ A-A stacking greatly increased fold change (Fig. S8C-D). These results suggest that 3’ A-A stacking preceding G108 also enhances strand displacement in the context of toehold-mediated strand displacement, extending our model of stacking-mediated branch migration enhancement.

In a complementary experiment, we added toehold base pairs to a length 7 polyA loop in either the A:U or U:A orientation, which revealed similar trends (Fig. S8F-L). Together, these data indicate that even in the context of toehold-mediated strand displacement, stacking interactions immediately preceding the invader domain can enhance branch migration and suppress dynamic range.

These results suggest that there are two separable mechanistic bases by which disrupting stacking within the 3’ end of the loop simultaneously alters sensitivity and fold change: i) disrupting stacking increases the rate of loop closure, shortening the ligand-binding window time and thus decreasing sensitivity; and ii) disrupting stacking decreases EP strand displacement efficiency, thus increasing fold change through increasing ON-state gene expression.

### Remote toehold EP architecture uncouples tradeoff between sensitivity and dynamic range

To evaluate whether the apparent tradeoff between sensitivity and fold change is an inherent relationship for the *Cbe pfl* EP architecture, or mainly a result of the polyA EP loop, we randomized the loop region and performed FACS-seq (38–40) to construct dose response curves for 3,289 length 7 nt loop variants (Fig. S9). Characterizing variants at this scale allowed us to construct a functional landscape mapping variants onto EC_50_ and fold change axes (Fig. S10A). This landscape revealed a tradeoff between sensitivity and dynamic range (Fig. S10A), suggesting that this tradeoff is inherent to the *Cbe pfl* EP loop-only architecture.

Based on this observation, we next sought to redesign the EP architecture to separate these two phenotypes by including a remote toehold. Remote toehold systems use a design concept from DNA nanotechnology consisting of a toehold connected to an invader by an extendible tether that slows down strand displacement by imposing an entropic cost on the invader:substrate duplex, allowing fine-tuning of strand displacement kinetics (52). We applied the remote toehold architecture in the context of the ZTP riboswitch by modifying the EP to consist of a synthetic toehold hairpin, which is separated from the AD and invader by variable length polyA tethers on both sides (Fig. 3A-B). Extending either one or both of these tethers should simultaneously impose two penalties: a kinetic penalty on the initial nucleation of P3 strand displacement, thus enhancing sensitivity, and an entropic penalty that slows down branch migration once initiated, enhancing dynamic range (50, 52).

We observed that extending these tethers on either or both sides results in dual enhancement of sensitivity and dynamic range (Fig. 3C-E). Increasing tether length enhances sensitivity as predicted (Fig. 3D), and fold change is increased by increasing ON-state especially for 5’ tether extensions, up to a point where increasing tether length causes baseline leak expression to increase (Fig. S11). This results in the remote toehold architecture showing a Pareto front-type of tradeoff with better sensitivity and fold-change than the loop variants characterized by FACS-seq (Fig. 3F, S10B).

These findings led us to conclude that the apparent tradeoff between sensitivity and dynamic range is not inherent to the ZTP riboswitch itself, but rather arises from the WT *Cbe pfl* EP loop-only architecture that has the same loop nucleotides influencing the initial EP nucleation attempts and the downstream branch migration process needed to form a complete terminator. Redesigning the EP architecture allowed us to achieve dual improvement in sensitivity and dynamic range that was precluded in the loop-only architecture.

### Natural ZTP riboswitch EPs use multiple approaches to enhance sensitivity

The amenability of *Cbe pfl* to drastic EP architectural changes motivated us to investigate whether natural ZTP riboswitch EPs contain architectures beyond the loop-only EP architecture common to the model ZTP switches previously characterized (10, 34, 35, 53). To do so, we bioinformatically identified all transcriptional ZTP riboswitches annotated in the Rfam entry RF01750 with a similar invader:P3 strand displacement fold as *Cbe pfl* (287 sequences; see **Methods**). Using secondary structure prediction of the spacer regions between P3 and the invader domain, we classified EPs into three architectural subtypes: 1) unstructured loop-only like *Cbe pfl* (158 sequences), 2) remote toehold EPs separated from the AD or invader by mismatches or internal loops (86 sequences), or 3) direct toehold EPs including 2 or more base pairs preceding the invader:P3 junction that could serve as toeholds of EP strand displacement (43 sequences) (Fig. 4A-B).

**Figure 4:**
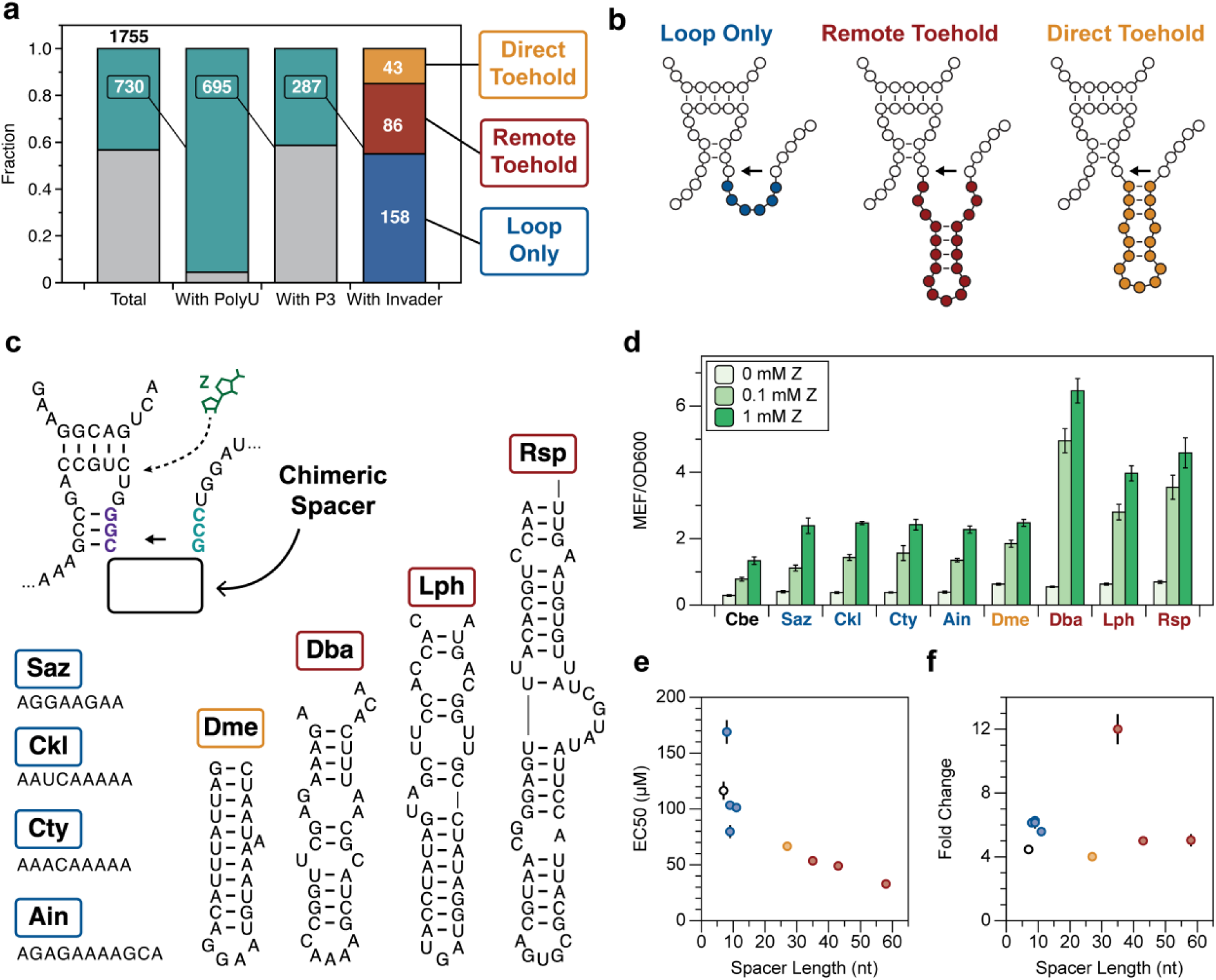
Natural ZTP riboswitch EPs use multiple approaches to enhance sensitivity. (a) The EPs of 1755 putative ZTP riboswitches were downselected based on the presence of a downstream polyU tract (730/1755), the presence of a consensus P3 stem (695/730), and the presence of an invader domain that pairs with P3 and competes with the pseudoknotted region for base pairs (287/695). These 287 EP candidates were manually classified into 3 categories based on the spacer region between P3 and the invader as indicated in panel (b): loop only, remote toehold, and direct toehold. (c) Secondary structure schematic of a panel of loop only, direct toehold, and remote toehold EPs (color coding as in panels (a) and (b) selected for further functional characterization. Illustrated regions were chimerically inserted in place of the *Cbe pfl* polyA terminator loop. The Rsp spacer region is a remote toehold because the invader includes a G108Δ deletion, so G108 was deleted from the invader region in the Rsp chimera. A table of the full sequences of these riboswitches and their genomic loci can be found in Table S1. (d) In vivo fluorescence data for chimeric riboswitches ordered according to spacer length, with Cbe indicating the WT *Cbe pfl* sequence. (e) EC_50_ measurements of chimeric ZTP riboswitches shown in (d). (f) Fold change of chimeric ZTP riboswitches shown in (e). Bars in panel (d) indicate average MEF/OD_600_ values over n = 9 replicates, with error bars indicating standard deviation. Data in (e) and (f) are determined as described in **Methods**.

Loop-only EPs were found to be similar in length to the *Cbe* loop on average (Fig. S12A), and biased toward being enriched for purines, with many loops possessing 100% purine content (Fig. S12B), suggesting that other ZTP riboswitches also employ stacking-mediated sensitivity enhancement like *Cbe pfl*. Direct toehold EPs are shorter in overall length and significantly enriched for AU content relative to remote toehold EPs, and rarely exceed 5 bp in length (Fig. S12C-E), suggesting that strong direct toeholds are disfavored relative to weaker toeholds. Additionally, we observed a strong bias in the orientation of direct toehold base pairs, with the three nearest pairs to the invasion site being significantly enriched for R-Y orientation relative to Y-R (Fig. S12F-G). Based on our observation that U-A toeholds result in very poor dynamic range relative to A-U toeholds (Fig. S8 C,H,J) we reasoned that these ZTP EPs are under purifying selection to avoid suppressing ON-state gene expression.

To test how these natural EP configurations contribute to riboswitch function, we constructed chimeric riboswitches by grafting the spacer regions from various natural sequences in place of the 7 nt polyA loop in *Cbe pfl*. We selected 4 loops longer than 7 nt, a relatively long direct toehold, and 3 remote toeholds of varying lengths (Fig. 4C). We observed that all of the spacer regions supported efficient switching by the chimeric riboswitches (Fig. 4D), and that the length of the spacer region was correlated with riboswitch sensitivity (Fig. 4E). As in the case of the synthetic remote toehold (Fig. 3), sensitivity enhancement by natural remote toeholds did not result in suppression of fold change (Fig. 4F), supporting our earlier conclusion that the remote toehold architecture uncouples the sensitivity-dynamic range tradeoff observed in the loop-only architecture. In fact, the Dba chimera has better sensitivity and fold change than any of the synthetic remote toehold variants.

Together, these analyses reveal that the principles uncovered by functional mutagenesis of the *Cbe pfl* ZTP riboswitch can help explain the sequence diversity of transcriptional ZTP riboswitch EPs, and that these natural riboswitches have evolved to exploit multiple EP architectural solutions to tune riboswitch sensitivity.

### Kinetics of EP nucleation also control translational ZTP riboswitch sensitivity

The remote toehold architecture reminded us of the layout of the *Pectobacterium carotovorum rhtB* riboswitch (*Pca*), a translational ZTP riboswitch previously characterized by Breaker and colleagues, which controls translation initiation by using the start codon of the *rhtB* gene as part of its EP invader domain (Fig. 5A) (53). Translational riboswitches are generally thought to operate as thermodynamic switches (27, 32), although growing evidence is suggesting that some riboswitches may control translation initiation in a kinetic regime (54). If true, such translational riboswitches should feature ligand-binding time windows, and therefore should be able to be sensitized by slowing EP nucleation. We therefore sought to test this prediction through functional characterization of *Pca* EP variants using the principles learned about tuning sensitivity from studying transcriptional ZTP riboswitches.

**Figure 5:**
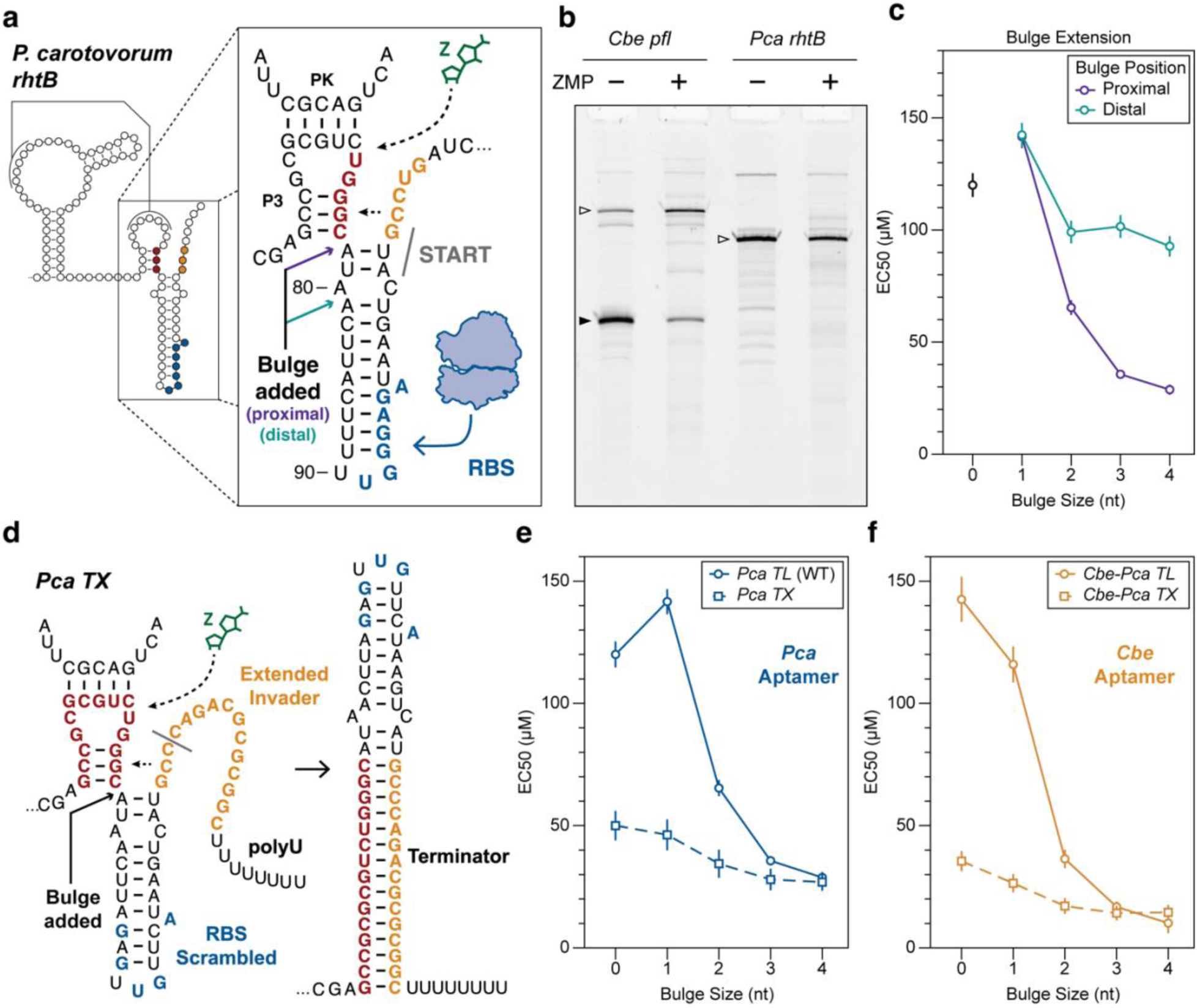
Kinetics of EP nucleation also control translational ZTP riboswitch sensitivity. (a) Schematic of the *Pca rhtB* ZTP riboswitch, which regulates translation initiation. In this mechanistic model based on findings of Breaker and colleagues (53), a ribosome binding site (RBS) is located within the distal portion of the EP, while the strand displacement switching mechanism is initiated by the G of the start codon as the invading nucleotide. Arrows indicate the positions where optional polyA bulges were inserted: proximal (purple, 77-A_n_-78) or distal (teal, 80-A_n_-81). (b) Single-round in vitro transcription reactions of the *Cbe pfl* and *Pca rhtB* riboswitches in the presence or absence of ZMP (1 mM). Filled triangles indicate the terminated length, and open triangles indicate full-length product based on the length of the linear template DNA. Source gel images of 2 replicates can be found in Fig. S14. (c) EC_50_ measurements of proximal and distal bulge insertion mutants. (d) Schematic of the synthetic EP designed to convert the *Pca rhtB* riboswitch to transcriptional regulation (*Pca TX*). The native RBS was scrambled, and the first 6 codons of the *rhtB* gene were replaced by a synthetic invader domain coupled to a polyU tract designed to form a functional intrinsic terminator. Arrow indicates the position where optional polyA bulges were inserted. (e) EC_50_ measurements of *Pca rhtB* (*Pca TL*) and *Pca TX* bulge variants. (f) EC_50_ measurements of chimeric riboswitches *Cbe-Pca TL* and *Cbe-Pca TX* which consisted of the EPs of *Pca TL* and *Pca TX* grafted onto the AD of *Cbe pfl*. Data in panels (c), (e), and (f) are determined as described in **Methods**.

We first confirmed that *Pca* does not regulate transcription elongation in a ligand-dependent fashion, like *Cbe pfl* or other transcriptional ZTP riboswitches, by showing that it does not function in an in vitro transcription reaction (Fig. 5B). We next tested the predicted switching mechanism of the *Pca* EP by mutating key regions, confirming their importance for switching function (Fig. S13A).

To investigate whether the kinetics of EP nucleation affect riboswitch sensitivity, we introduced polyA bulges near the invader:P3 junction to slow the formation of the predicted start codon-sequestering structure. These polyA bulges were either placed proximal to the strand displacement initiation site, to convert the EP to a remote toehold architecture, or 1 nt distal, to extend the A80:C103 mismatch into an internal loop (Fig. 5A). We observed that proximal polyA bulge extension enhanced sensitivity in a manner similar to remote toehold leash extension, while distal bulge extension also enhanced sensitivity though to a lesser extent (Fig. 5C). As expected, these bulge extension series, which disrupt the region responsible for mediating switching function (Fig. S13A), resulted in length-dependent decreases in fold change caused by increases in riboswitch leak (Fig. S15A-C).

We next sought to confirm that these same polyA bulges would enhance sensitivity in a transcriptional version of this riboswitch that should operate in a kinetic regime. To do so, we converted the *Pca* EP to a transcriptional regulatory mechanism *(Pca TX*) by scrambling the native RBS, extending the native invader domain to fully complement P3, and replacing the 9 codons of the *rhtB* gene with a polyU tract followed by a synthetic RBS (Fig. 4D). We confirmed by single round IVT that this modified riboswitch functions by regulating transcription termination (Fig. S13B). Interestingly, we found that the *Pca TX* variant was more sensitive than the wildtype translational version, and importantly was also further sensitized by the insertion of a polyA bulge on the 5’ side of the toehold (Fig. 4E).

To ensure that no unknown features of the *Pca* AD were confounding our mutagenesis approach, we constructed chimeric versions of these riboswitches with the *Pca* AD replaced by the distantly-related *Cbe pfl* AD to create translational (*Cbe-Pca TL*) and transcriptional (*Cbe-Pca TX*) versions, and observed very similar sensitization patterns (Fig. 4F, S13B). Interestingly, for both *Pca* and *Cbe pfl* ADs, their respective transcriptional and translational versions approach the same EC_50_ value as the polyA bulge is extended, suggesting that as EP nucleation is slowed, EC_50_ values approach a common minimum value that does not rely on EP mechanism. All four of these mutational series resulted in expected length-dependent increases in riboswitch leak leading to decreases in fold change (Fig. S15D-I).

Together, these results demonstrate that the sensitivity of translational ZTP riboswitches can be enhanced through EP architecture changes that slow EP nucleation kinetics, generalizing our understanding of kinetically driven riboswitches to include examples of translational regulation.

### Delaying EP nucleation sensitizes diverse riboswitches

Finally, we sought to apply our understanding of how EP nucleation kinetics tunes riboswitch sensitivity to other riboswitch classes which feature diverse aptamer structures and direct toehold EP architectures, including riboswitches for purines and fluoride.

The model *B. subtilis pbuE* riboswitch senses adenine through an AD consisting of a three-way junction, and features a weak toehold connecting the AD and invader (Fig. 6A). We created a remote toehold architecture by inserting a polyA bulge at the strand displacement initiation site (Fig. 6A), and observed that polyA bulge length correlated with enhanced riboswitch sensitivity and fold change in a manner reminiscent of the remote toehold extensions in ZTP (Fig. 6B, Fig. S16A-B). Importantly, these polyA insertions disrupt a polyU tract previously reported to be an important regulator of sensitivity and dynamic range via putative RNAP pausing (55, 56), although we observe improved sensitivity and dynamic range for all polyA bulge variants relative to WT *pbuE* (Fig. 6B, S16A-B).

**Figure 6:**
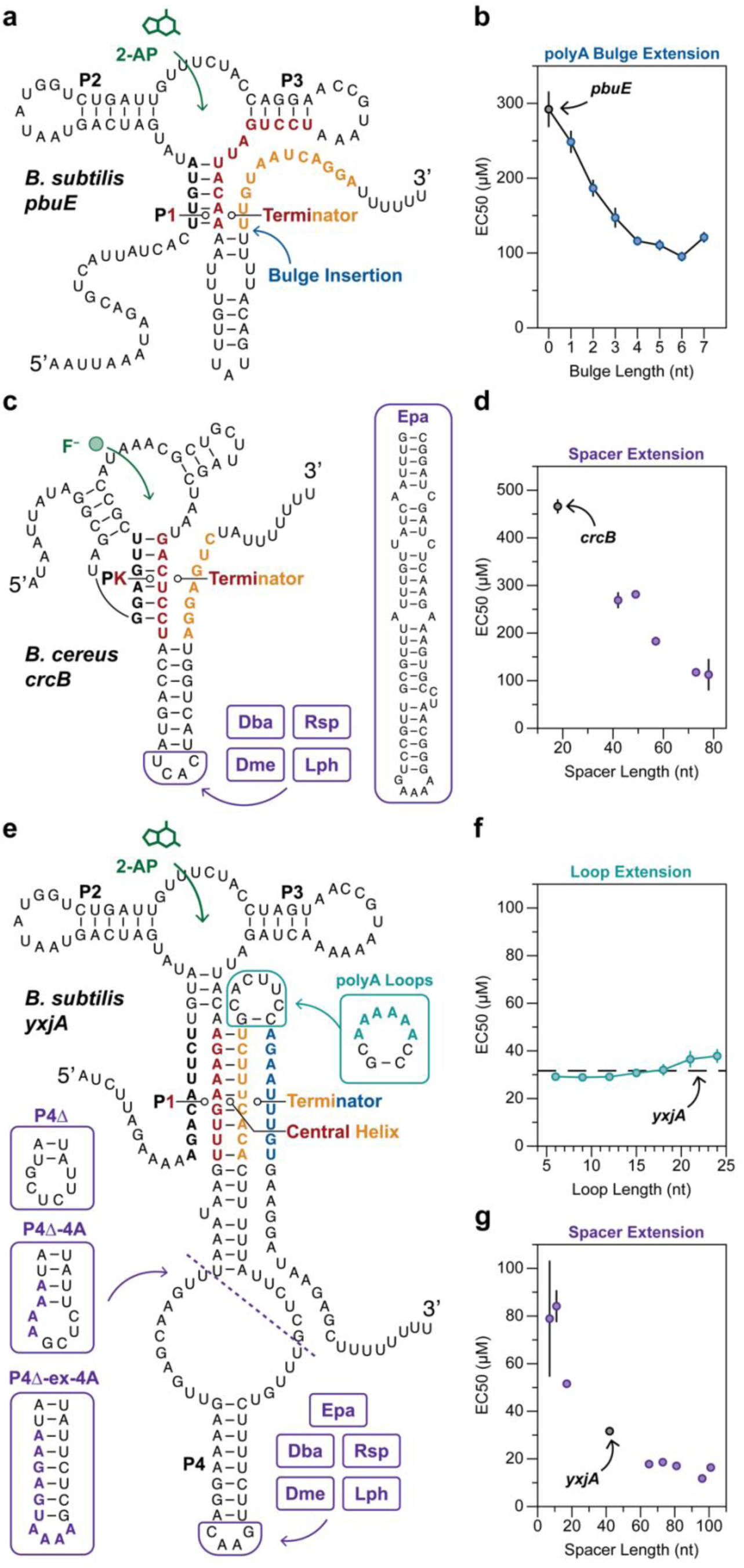
Delaying EP nucleation sensitizes diverse riboswitches. (a) Schematic of the *Bsu pbuE\** leader truncation variant of the *pbuE* transcriptional ON riboswitch (55). A weak direct toehold facilitates nucleation of the EP, in which an invader (gold) strand displaces P1 (black and red) to form the terminator hairpin. Arrow indicates the position of polyA bulge insertion (b) EC_50_ measurements of *pbuE* polyA bulge variants. (c) Schematic of the *Bce crcB* fluoride transcriptional ON riboswitch. A direct toehold facilitates nucleation of the EP, in which an invader (gold) strand displaces the pseudoknot (black and red) to form the terminator hairpin. Boxes show terminator extension inserts appended to the end of the terminator hairpin. (d) EC_50_ measurements of *crcB* terminator extension variants. (e) Schematic of the *Bsu yxjA* transcriptional OFF riboswitch. A weak toehold comprising P4 and an internal loop facilitates nucleation of the EP, in which an invader (gold) strand displaces P1 (black and red) to form the central helix. The terminator loop (teal) can close to allow a second invader (blue) to strand displace the central helix (red and gold) to form the terminator hairpin. The second strand displacement process gives this switch OFF logic. Boxes show terminator loop extension variants, and two types of P4 spacer mutations: P4 truncations with different lengths added back, and P4 extensions. (f) EC_50_ values of *yxjA* terminator loop extension mutants. (g) EC_50_ measurements of *yxjA* P4 spacer mutations. Data in panels (b), (d), (f), and (g) are determined as described in **Methods**.

The *Bacillus cereus crcB* fluoride riboswitch uses an AD consisting of an H-type pseudoknot, and features a 7 bp direct toehold spacer region (Fig. 6C). We extended this direct toehold by chimerically grafting on the long ZTP spacer regions characterized in Fig. 4C, as well as a long (63 nt) spacer region from the *Enterococcus pallens eno* (Epa) fluoride riboswitch identified using similar bioinformatic analysis as described above (Fig. 6C, **Methods**). These spacer region extensions substantially sensitized the *crcB* fluoride riboswitch in a length-dependent manner (Fig. 6D), resulting in commensurate length-dependent decreases in fold change corresponding to increasing riboswitch leak (Fig. S16C-D).

We next sought to apply our findings to transcriptional riboswitches that feature an OFF regulatory logic. In previous work, we developed a synthetic OFF variant of the *Cbe pfl* ZTP riboswitch by inserting an 8 nt ‘flipping domain’ that inverts the EP regulatory logic (Fig. S17A) (31). To test if this OFF-switch could also be sensitized by delaying EP nucleation, we extended the polyA loop and observed length-dependent sensitization similar to what was observed for the WT ON-switch (Fig. S17B-C).

Finally, we investigated the *B. subtilis yxjA* purine riboswitch, which uses an AD similar to *pbuE* and which previous work demonstrated to possess two competing strand exchange processes (Fig. 6E) (57). We wondered if, like *pbuE*, the nucleation of the first strand displacement process (central helix) determines the ligand-binding time window, or if rather the nucleation of the terminator hairpin is the ligand-binding window time-determining step. To interrogate this question, we first varied the length of the terminator hairpin from 4A to 22A, which had a strong negative effect on dynamic range (Fig. S16E-F), but had little effect on sensitivity (Fig. 6F). Next, we manipulated the long remote toehold that nucleates the central helix. We either shortened this region by deleting the 5’ tether and the P4 stem, or we extended this region by grafting the spacers from ZTP and fluoride riboswitch EPs onto the end of P4. We observed that the shortened P4Δ mutants became less sensitive than wild type, and the extended variants all became more sensitive, with EC_50_ correlating with the overall length of the spacer region (Fig. 6G, S16G-H). These results confirm that the initial strand displacement step that forms the central helix determines the ligand-binding time window, as in *pbuE*, and that the second strand displacement step, which allows the terminator to form, occurs after the riboswitch has committed to produce a given regulatory outcome.

Taken together, these results demonstrate that the principle of tuning riboswitch sensitivity through EP sequence/structural changes that control EP nucleation kinetics are generalizable, and can be used to sensitize any riboswitch that operates in a kinetic regime.

### Minimum achievable EC_50_ from in vivo kinetic riboswitches differs from in vitro aptamer:ligand K_D_ values

Early discoveries in the riboswitch field demonstrated a large discrepancy between in vitro AD:ligand K_D_ measurements and transcriptional riboswitch EC_50_ measurements, leading to the general acceptance of the model that EC_50_ is elevated in these riboswitches because the kinetic constraint of transcription prevents the AD from reaching equilibrium with its ligand (28). Having discovered and developed a set of approaches to extend the AD:ligand interaction time and sensitize various riboswitch classes, we asked whether these enhancements in sensitivity allow EC_50_ to approach AD:ligand K_D_ as interaction time increases. Interestingly, for the various methods of sensitization we employed, EC_50_ seems to asymptotically approach some minimum value for each riboswitch class as the kinetic barrier to EP nucleation increases (Fig. 1B, 3D, 4E, 5E-F, 6B, 6D, 6G, S17C). However, plotting the minimum achievable EC_50_ values in comparison to experimental K_D_ values reveals a remaining 2-200x discrepancy (Fig. 7). To ensure that this discrepancy was not caused by membrane effects limiting intracellular ligand concentration in our in vivo assay, we compared our in vivo EC_50_ measurements to literature values for in vitro EC_50_, which showed that WT *Cbe pfl* and *crcB* replicate in vitro EC_50_ values almost exactly (Fig. 7A,D). However, the observed in vivo EC_50_ for *pbuE* exceeds the in vitro EC_50_ by ∼10x (Fig. 7B), indicating that membrane permeability may confound our in vivo EC_50_ measurements for 2AP-sensing riboswitches.

**Figure 7:**
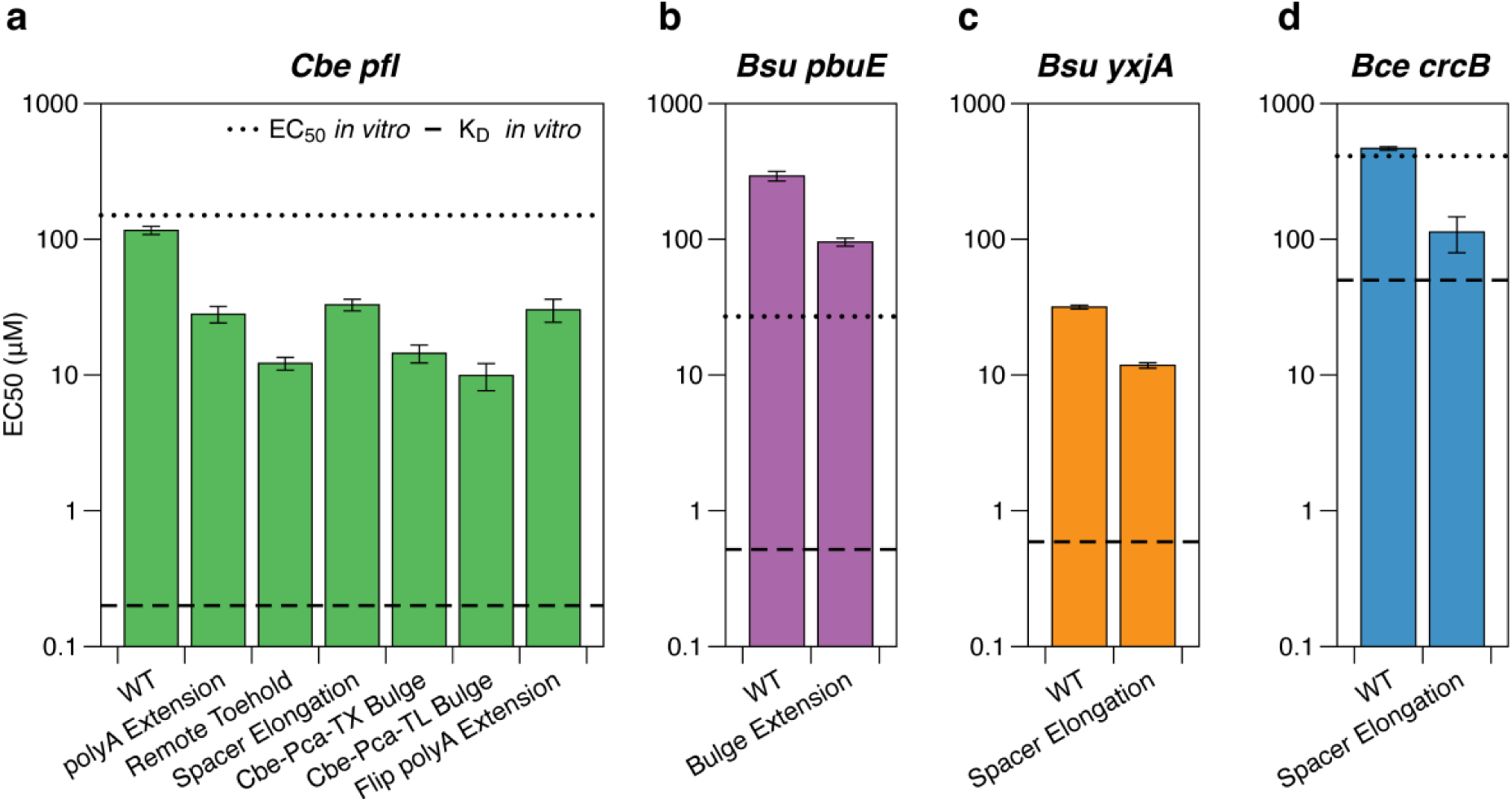
Minimum achievable EC_50_ from in vivo kinetic riboswitches differs from in vitro aptamer:ligand K_D_ values. (a) Cbe *pfl* ZTP riboswitch EC_50_ plotted with the minimum EC_50_ observed by various sensitization approaches. Dotted line indicates average of two literature values for in vitro EC_50_ obtained in different transcription conditions (35, 53). Dashed line indicates AD K_D_ for ZMP (53). (b) *Bsu pbuE* 2AP-binding riboswitch EC_50_ plotted with the minimum EC_50_ observed by polyA bulge extension. Dotted line indicates in vitro 2AP EC_50_ (60), and dashed line indicates AD K_D_ for 2AP (27). (c) *Bsu* yxjA 2AP-binding riboswitch EC_50_ plotted with the minimum EC_50_ observed by spacer elongation. No research group has successfully determined in vitro EC_50_ (61). Dashed line indicates AD K_D_ for 2AP (62). (d) *Bce crcB* fluoride riboswitch EC_50_ plotted with the minimum EC_50_ observed by spacer elongation. Dotted line indicates in vitro EC_50_ (63), and dashed line indicates AD K_D_ for fluoride inferred from similar aptamers in (64). Data in (a-d) are determined as described in **Methods**.

Focusing our attention on *Cbe pfl* and *crcB*, we hypothesized that minimum EC_50_ may be elevated relative to in vitro K_D_ because the cotranscriptional ensembles of these riboswitches feature abundant kinetically trapped states only capable of forming partial ligand contacts within the timeframe of transcription. An immediate upshot of this model is that aptamers with more extensive ligand interactions will have more possible partially bound states of lower affinity than the refolded structure, resulting in a larger discrepancy between cotranscriptional sensitivity and K_D_. This model is consistent with the observed trend that for *crcB,* which features a relatively simple ligand-binding interactions (58), only a ∼2x minimum EC_50_:K_D_ ratio is observed, while for ZTP riboswitches, which feature more numerous and elaborate ligand-binding interactions (59), a ∼50x minimum EC_50_:K_D_ ratio is observed (Fig. 7A). These findings emphasize the importance of other factors in limiting transcriptional riboswitch sensitivity besides the kinetic constraint of ligand-binding windows.

## DISCUSSION

Riboswitches are highly conserved gene regulatory elements controlling housekeeping and emergency-response functions in bacteria. Because of their key homeostatic roles, riboswitches must be finely tuned to allow the commensurate amount of gene expression in response to the intracellular concentrations of their cognate ligands. The demanding sensitivity and specificity requirements of ADs seem to be why they are among the most highly conserved RNA domains in all of biology (65). This high conservation of ADs could explain the observed natural diversity of EPs. As the more evolutionarily malleable riboswitch domain, EPs enable nearly invariant ADs to regulate entirely different steps of gene expression, and allow ADs with nM affinity to operate in the μM or mM sensitivity ranges (66, 67). However, the mechanisms by which EP sequence changes tune riboswitch function remain relatively unexplored.

We previously observed that the EP of the *Cbe pfl* ZTP riboswitch features an invader domain that can quantitatively reprogram riboswitch dynamic range by altering the kinetics of internal strand displacement (31). Other work has shown that the length of the *pbuE* EP toehold can tune dynamic range (55), and that nuanced EP sequence features can tune the sensitivity and dynamic range of the *Pseudomonas aeruginosa* guanidine-II translational riboswitch (68). In this work, we identify EP nucleation as a key determinant of riboswitch sensitivity in *Cbe pfl*, and describe three unique approaches used by natural ZTP riboswitches to delay EP nucleation and tune sensitivity: purine-rich loops, remote toeholds, and long spacer regions. Together, these findings are forming a complete sequence-structure-function understanding of each component of the ZTP riboswitch EP (Fig. S18), which reveals the outsized role of EP folding in determining all aspects of riboswitch function (67).

Importantly, we have shown that principles for tuning riboswitch sensitivity and fold change through manipulating EP folding pathways are general across diverse riboswitches that control different aspects of gene expression. Specifically, we characterized a translational ZTP riboswitch, *Pca rhtB*, and showed that it is sensitized by the introduction of kinetic barriers to EP nucleation, indicating that this riboswitch operates in a kinetic regime. This finding supports other work showing that some translational riboswitches are kinetically controlled, and are not all thermodynamic switches (54). While it is possible that transcription-translation coupling may contribute to the observed kinetic constraint for ligand binding for *Pca* (69), we did observe similar sensitization patterns when the EP was converted into a transcriptional mechanism. In principle, any translational switch could be forced to operate in a kinetic regime if the EP-nucleated state includes a kinetic trap that blocks re-formation of the ligand-binding competent AD structure, as is true in the case of ZTP riboswitches (34). For thermodynamically-operated translational riboswitches, ready interchange between ON and OFF states will be possible even in the presence of the EP, as is true in the case of adenine translational riboswitches (27, 70).

We extended our findings from the ZTP system to a panel of diverse riboswitches with different ADs and different regulatory logic (ON vs OFF), and show that kinetic barriers that delay EP nucleation result in riboswitch sensitization in all of these cases, demonstrating the generality of our findings. A deeper comparison of observed sensitivity improvements across these diverse riboswitches raises an intriguing question – is there a fundamental limit to the sensitivity of a kinetically driven riboswitch? While we were able to achieve 2.5-12-fold enhancements in sensitivity for various riboswitches, for most classes the EC_50_ values of the best performing variants asymptotically approach levels that remain 1-2 orders of magnitude greater than published AD:ligand K_D_ values (Fig. 7). Moreover, very similar minimum EC_50_ values are obtained for kinetically driven transcriptional and translation riboswitches that use the same AD (Fig. 5E,F), suggesting that minimum EC_50_ is a property set by the binding interaction between the ligand and the AD, but is also fundamentally limited by the constraints of a kinetically driven process. This could be due to two reasons: (1) There is a limit to the ability to extend the ligand-binding window time, perhaps due to the fundamental instability of kinetic transient out of equilibrium RNA folds, and thus there is no chance to ever approach the equilibrium AD:ligand interactions observed in K_D_ measurements of refolded RNAs; or (2) The AD ligand-binding interactions that occur during the out-of-equilibrium transient cotranscriptional RNA folding processes of kinetically driven switches are fundamentally different, with lower affinity than those characterized in vitro with refolded RNA. If the latter model, originally proposed by (28) is true, it could give insights into what is needed to develop drugs that target transient cellular RNA structures (71).

Conceptually, kinetically-operated riboswitches function more like dimmer switches or fuses than digital switches (28, 72). This study highlights the key role of RNA folding kinetics, especially those of EPs, in governing the functional properties of riboswitches in terms of sensitivity and dynamic range. Taken together with many discoveries characterizing the nuanced ways in which transcriptional dynamics can reciprocally control RNA folding (26, 28, 29, 63, 73–75), these results emphasize how natural systems exploit the biophysical properties of cotranscriptional RNA folding to generate riboswitch variants that meet the needs of diverse genetic contexts (37). This work also begins to generalize our understanding of riboswitch tuning strategies across diverse classes and across different modes of regulation. We anticipate these studies to guide further development of riboswitch-based biotechnologies (17–21), RNA-targeting drugs including antibiotics (4–7), and add to our general understanding of how out-of-equilibrium RNA folding dynamics impacts RNA function.

## ACKNOWLEDGMENTS

We thank John Marko and Petr Šulc for helpful discussions about polymer theory and RNA folding dynamics. We thank Yueyi Li for providing a fluorescein standard calibration for the microplate reader. We thank the Robert H. Lurie Comprehensive Cancer Center of Northwestern University in Chicago, IL, for the use of the Flow Cytometry Core Facility, which provided use of the BD Melody FACS sorter. Support for this work was provided by the National Institutes of Health grant R01GM130901 to J.B.L., the National Institutes of Health Training Grant T32GM008382 through the Northwestern University Molecular Biophysics Training Program to D.Z.B., the National Science Foundation Grant 2310382 to J. B. L., and the Northwestern University Rappaport Award for Research Excellence to D. Z. B. The content is solely the responsibility of the authors and does not necessarily represent the official views of the NIH or NSF.

## AUTHOR CONTRIBUTIONS

Conceptualization, D.Z.B and J.B.L.; Methodology, D.Z.B and J.B.L.; Formal analysis, D.Z.B and J.F., Investigation, D.Z.B and J.F.; Writing – Original Draft, D.Z.B. and J.B.L.; Writing – Review & Editing, D.Z.B., J.F., and J.B.L.; Visualization, D.Z.B.; Software, D.Z.B and J.F.; Supervision, J.B.L.; Funding Acquisition, J.B.L.

## RESOURCE AVAILABILITY

### Lead contact

Further information and requests for resources and reagents should be directed to and will be fulfilled by the Lead Contact, Julius Lucks.

### Materials availability

Reporter plasmids for select variants from each riboswitch class investigated in this study will be available through Addgene. See Supplementary Data File 1 for complete sequence information for all variants and Addgene IDs for select variants.

### Data and code availability

All fluorescence data is available in Supplementary Data File 1. The accession number for FACS-seq data reported in this paper is SRA: PRJNA1087340. Flow cytometry data will be deposited in FlowRepository pending server maintenance. All code is available on Github repository https://github.com/LucksLab/Bushhouse_Riboswitch_Sensitivity_2024.

