## Supplemental Information for "RNA folding kinetics control riboswitch sensitivity in vivo"

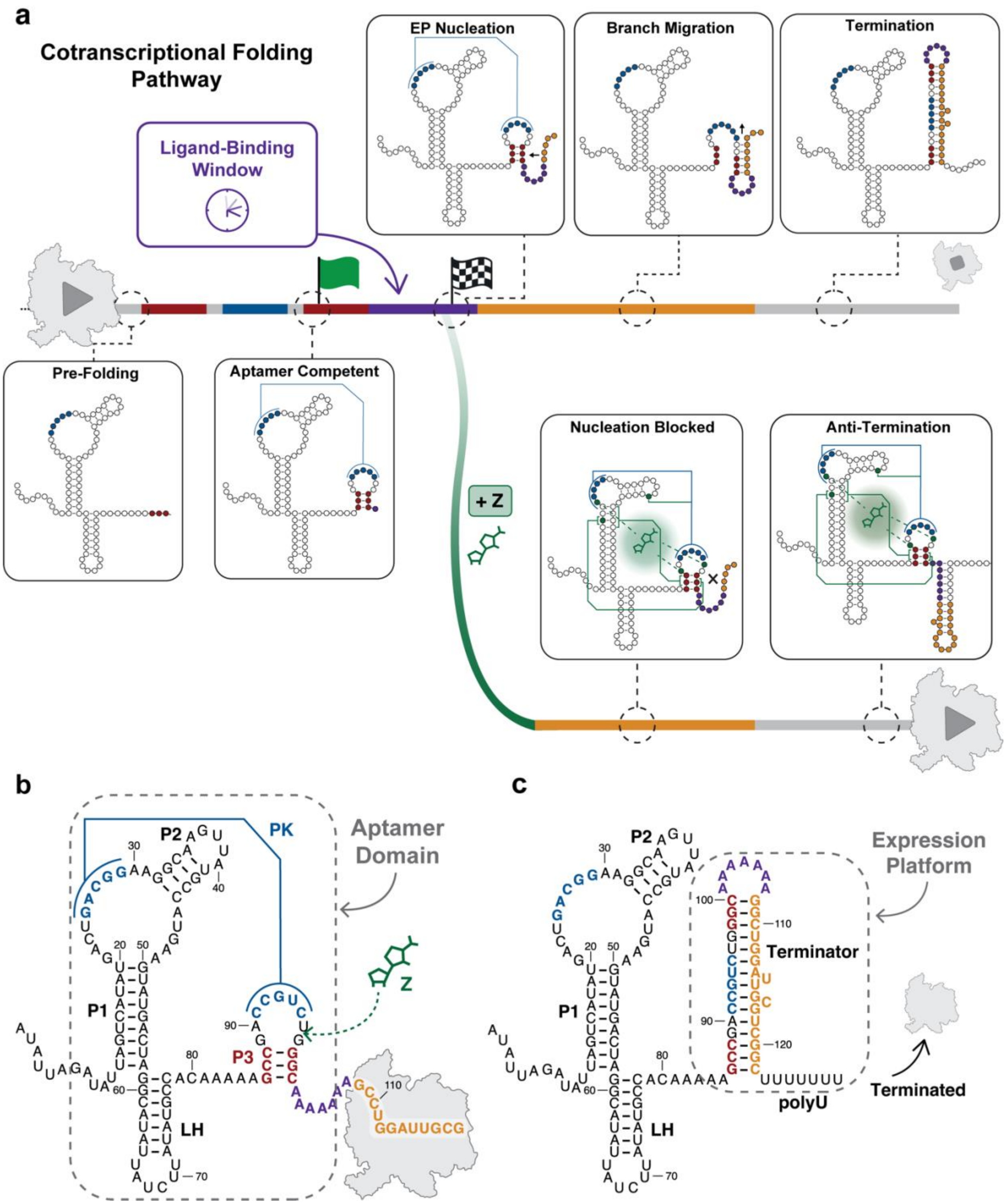


**Figure S1: *Cbe pfl* ZTP riboswitch folding pathway and secondary structure.** (a) Schematic of the *Cbe pfl* cotranscriptional folding pathway. After transcription of the full aptamer sequence, the ligand-binding AD domain quickly folds into the pseudoknoted (blue) competent structure (1–3). Ligand binding remains possible during the transcription of the downstream 7-10 nt (Ligand Binding Window, beginning with green start flag), until nucleation of the EP disrupts the AD (checkered flag) (1, 3). By default, EP nucleation initiates strand displacement branch migration that results in the formation of an intrinsic terminator that terminates transcription. However, if Z/ZMP/ZTP (green) is bound during the ligand-binding window, the riboswitch is diverted onto an alternative folding pathway in which EP nucleation is less likely to occur. Stabilization of the AD by ligand-binding interactions makes strand displacement less likely, leading to an increased frequency of anti-termination. The ligand-binding window can kinetically constrain riboswitch sensitivity (4). (b) Secondary structure of the ligand-binding competent *Cbe pfl* during transcription (3). The AD (dashed box) features a helix-junction-helix motif containing P1 and P2 pseudoknotted to the small P3 stemloop, connected by a non-conserved linker hairpin (LH). Binding of Z stabilizes P3 through numerous tertiary structural interactions (5, 6) (c) Secondary structure of terminated *Cbe pfl* (3). The EP (dashed box) comprises the terminator loop (purple) and invader domain (gold), which strand displaces all of P3, including the pseudoknotted region. RNA polymerase schematic depicted in gray.

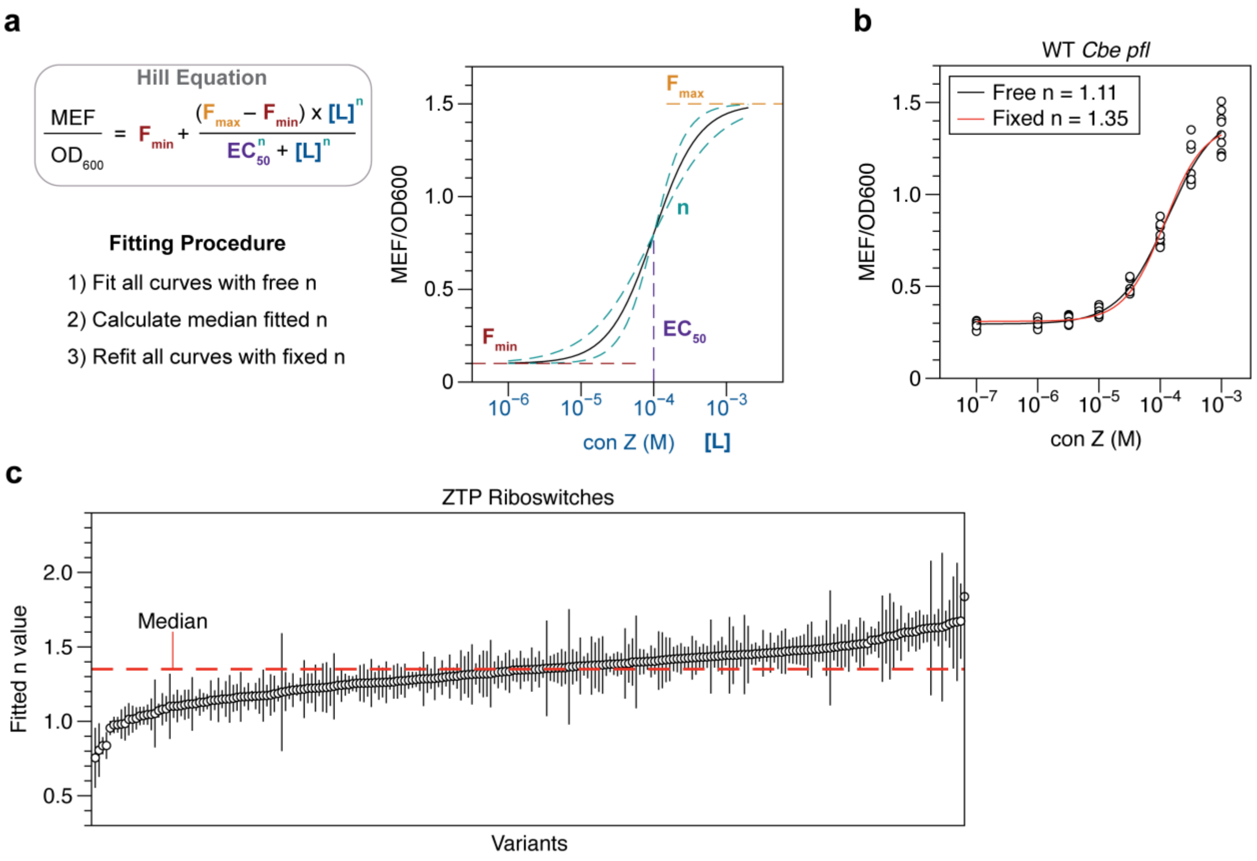


**Figure S2: Fitting procedure to determine dose-response characteristics of ZTP riboswitches.** (a) Dose response curves were obtained by fitting the Hill equation to all 72 measurements. Initially, all riboswitches of a given class were fit using an unconstrained Hill coefficient (b). The median value of all fitted n values (c) was used to refit all dose response curves, ensuring that all error for the fit would be captured in the $EC_{50}$ and fold change standard error. Error bars in panel (c) indicate standard error of the fit parameter n. See Supplementary Data File 1 for all fluorescence measurements and fits.

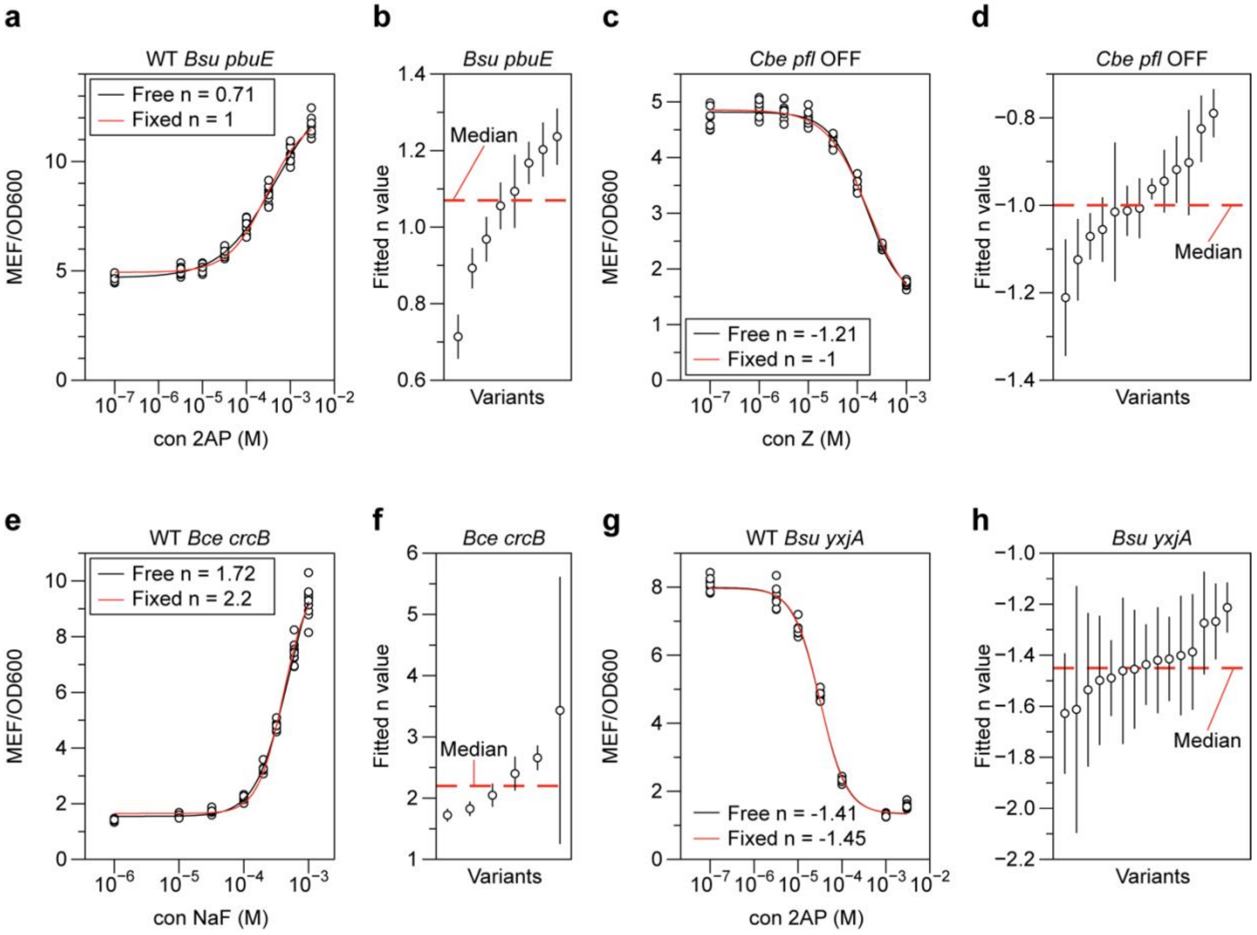


**Figure S3: Fitting procedure to determine dose-response characteristics of various riboswitches.** Dose response curves were obtained by fitting the Hill equation to all 72 measurements. Initially, all riboswitches of a given class were fit using an unconstrained Hill coefficient (a), (c), (e), (g). The median value of all fitted n values for a given riboswitch class (b), (d), (f), (h) was used to refit all dose response curves, ensuring that all error for the fit would be captured in the $EC_{50}$ and fold change standard error. Error bars in panels (b), (d), (f), and (h) indicate standard error of the fit parameter.

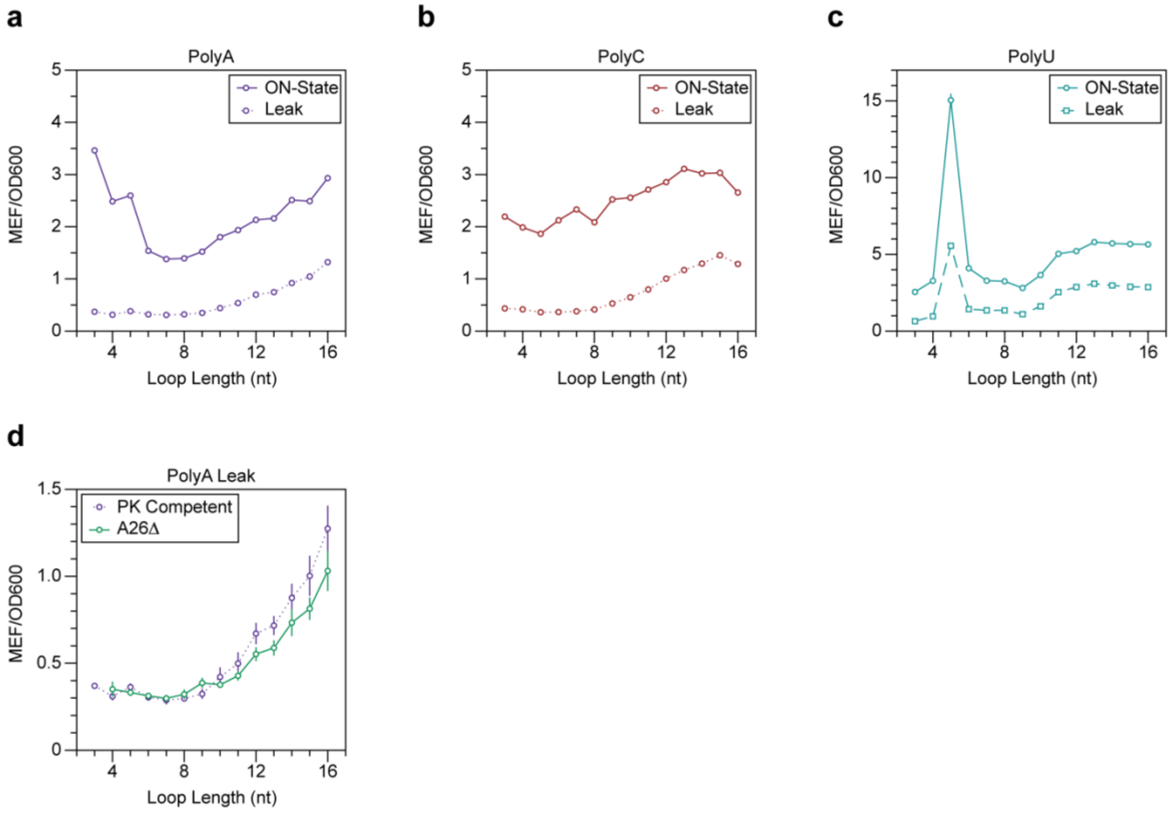


**Figure S4: Increasing loop length affects riboswitch leak by decreasing termination efficiency.** ON-state ($F_{max}$) and leak ($F_{min}$) for (a) polyA loop length variants, (b) polyC loop length variants, and (c) polyU loop length variants. (d) Gene expression at 0 mM Z for polyA loops with and without the A26Δ mutation. Data are determined as described in **Methods**.

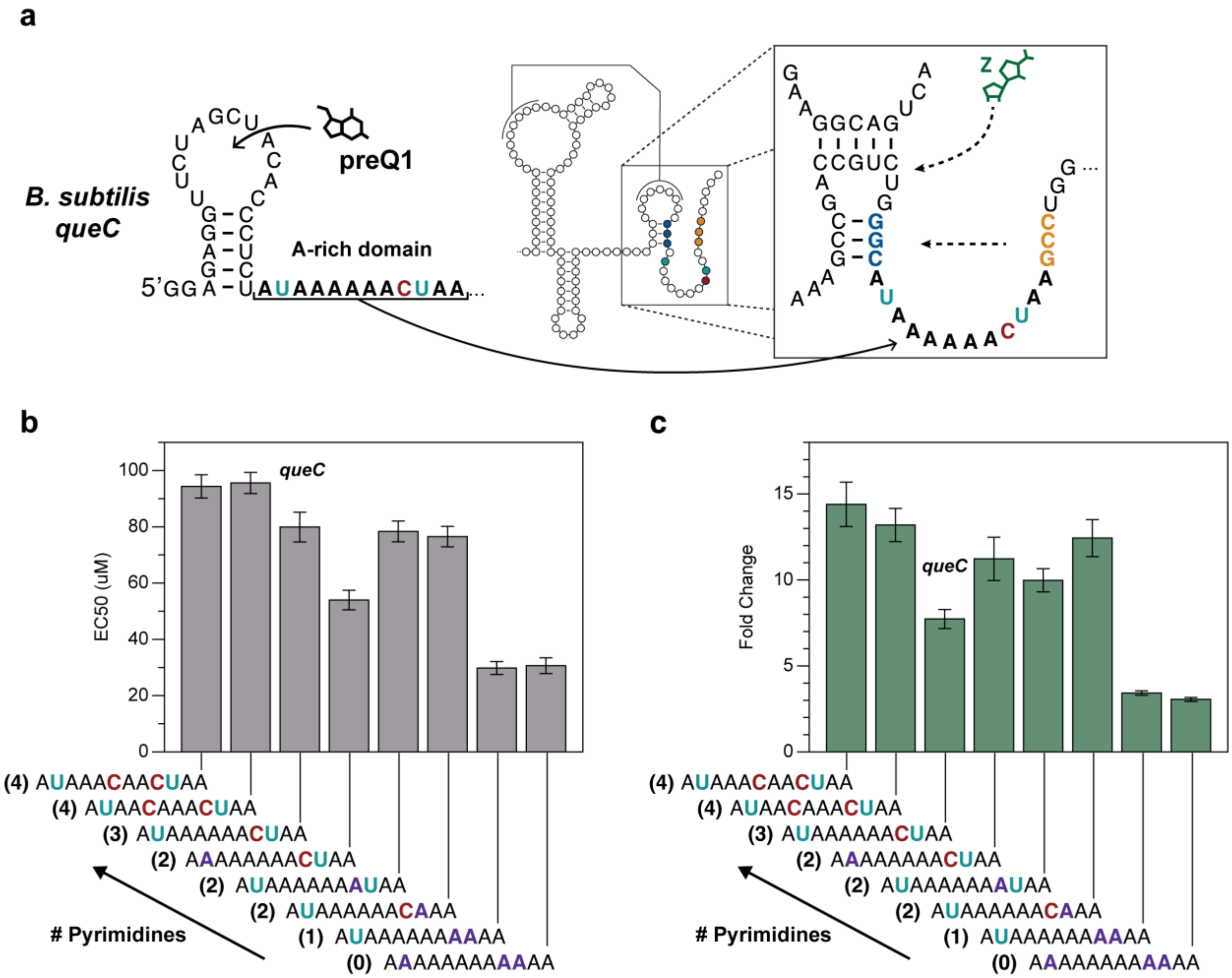


**Figure S5: Disruptions to A-A stacking in a model A-rich sequence recapitulate functional tradeoff between sensitivity and fold change in a ZTP riboswitch EP.** (a) Schematic detailing how the A-rich single stranded region from the *Bsu queC* preQ1 riboswitch was chimerically inserted in place of the *Cbe pfl* polyA terminator loop. Additional mutations were made to this chimeric loop to interrogate the role of stacking within this region in mediating sensitivity and fold change. Past work by Eichhorn *et al.* on the *Bsu queC* A-rich domain showed with NMR that A-A stacking interactions rigidify this domain, and that the introduction of a single A→C mutation in the A-rich core locally destabilizes the stacking interactions in this domain (7). We chimerically replaced the 7A loop of *Cbe pfl* with this 12 nt A-rich domain from *Bsu queC*, characterized dose response curved and extracted (b) $EC_{50}$ and (c) fold change values for *queC* loop variants with different numbers of pyrimidine residues. Compared with a perfect polyA loop of the same length, the *Bsu queC* domain, which contains three pyrimidine residues, including two near the 3' end, resulted in significantly worse sensitivity but higher fold change. Introducing additional A→C mutations into the A-rich core resulted in further desensitization of chimeric riboswitch mutants relative to the WT *Bsu queC* domain, and simultaneous correction of both 3'-proximal pyrimidines to A was required to restore sensitivity to the same level as polyA. Data in panels (b) and (c) are determined as described in **Methods**.

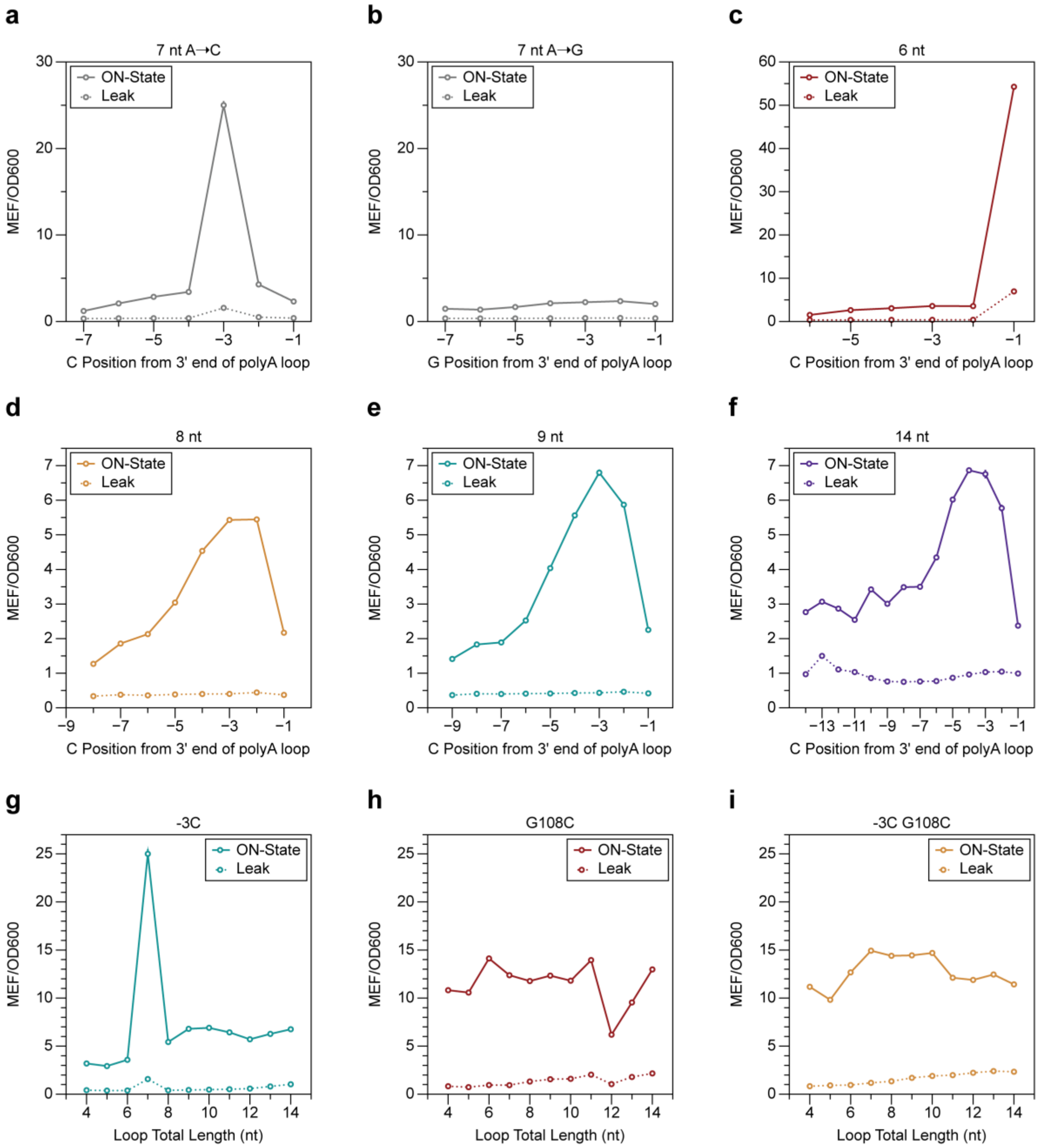


**Figure S6: Disruptions to loop stacking mediate large increases in ON-state gene expression.** ON-state ($F_{max}$) and leak ($F_{min}$) for EP variants of the *Cbe pfl* ZTP riboswitch (see Figure 2): (a) polyA length 7 nt A→C mutational scan, (b) polyA length 7 nt A→G mutational scan, (c) polyA length 6 nt A→C mutational scan, (d) polyA length 8 nt A→C mutational scan, (e) polyA length 9 nt A→C mutational scan, (f) polyA length 14 nt A→C mutational scan, (g) varying polyA loop lengths where the third to last residue relative to the 3' end of the loop has been mutated to C (-3C), (h) varying polyA loop lengths where the initial invading residue G108 has been mutated to C (G108C); and (i) varying polyA loop lengths with both the -3C and G108C mutations. Data are determined as described in **Methods**.

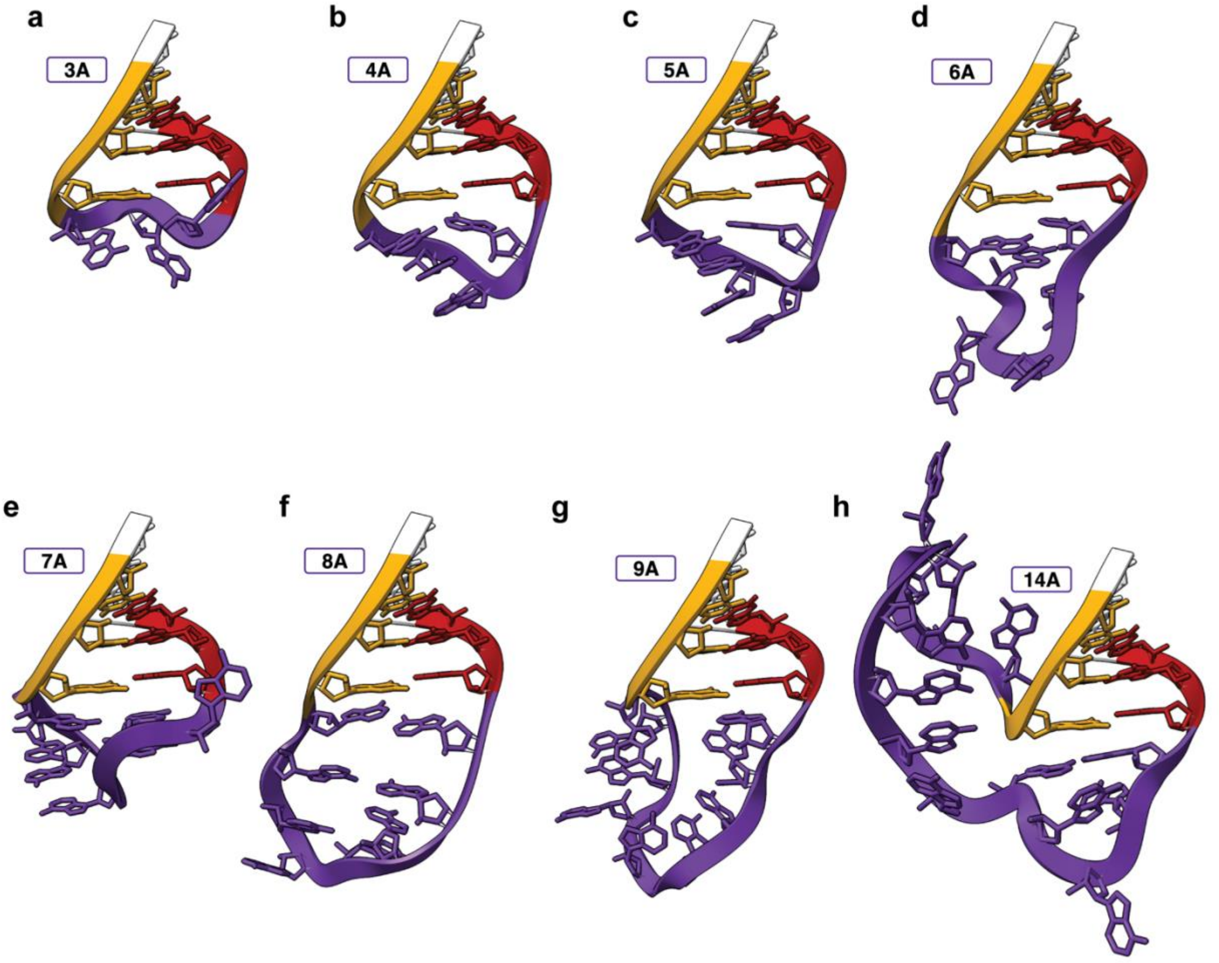


**Figure S7: Vfold3D modeling of nucleated polyA loops.** (a-h) Vfold3D models of nucleated terminator hairpins with different length polyA loops. The substrate strand of P3 (red) and the invader (gold) form a duplex by closing the polyA loop (purple). Loops shorter than 6A do not appear capable of aligning parallel rings, and therefore should not be able to form a 3’ stacking platform to stabilize G108 (See Figure 2F).

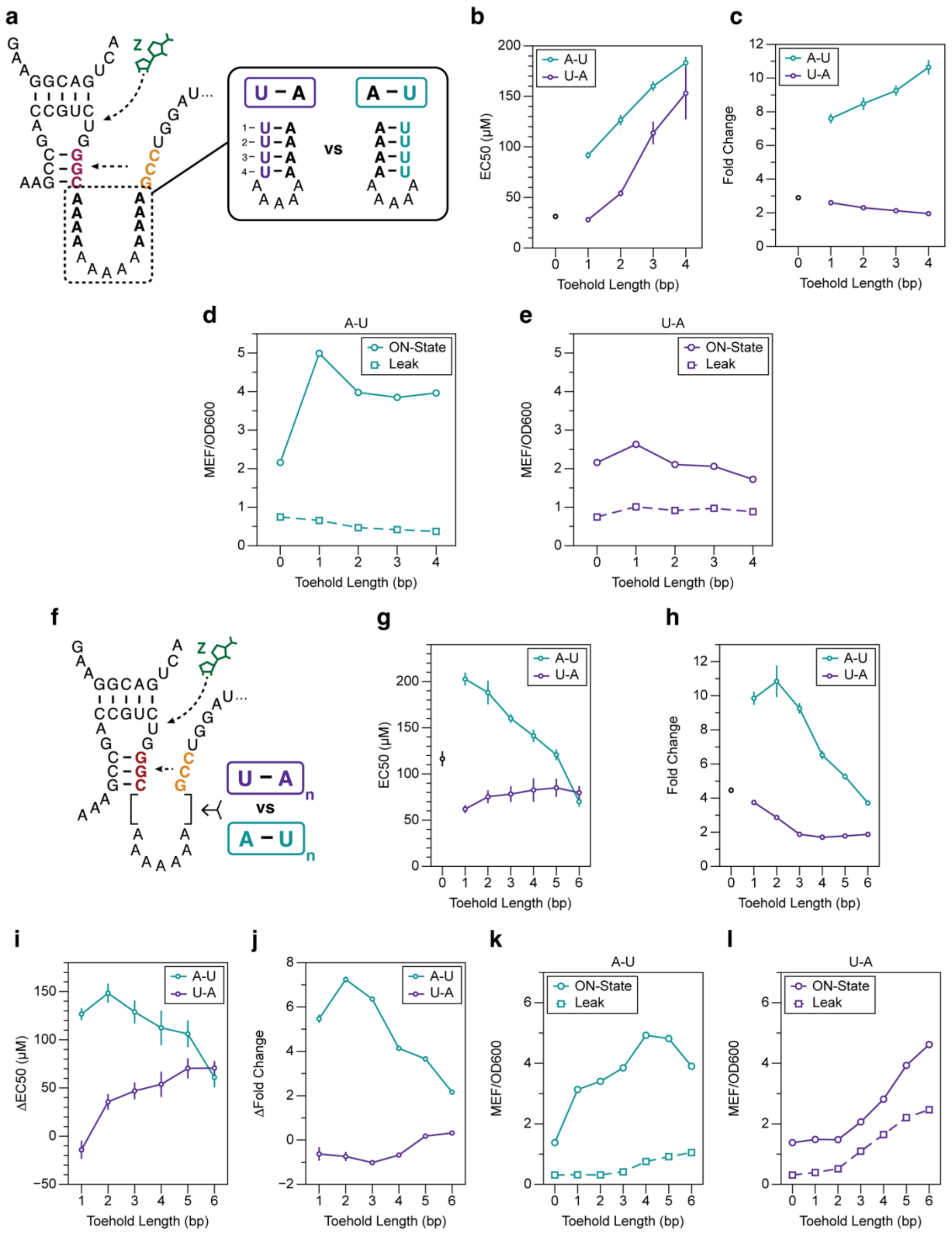


**Figure S8: 3' stacking within strand displacement toehold controls riboswitch dynamic range.** (a) Schematic detailing how a length 13 polyA loop was progressively shortened by the introduction of A→U mutations on either the 5’ side of the loop (purple) or the 3’ side of the loop (teal). The introduction of each additional A→U mutation converts the complementary A residue on the other side of the loop into a toehold base pair and shortens the remaining polyA loop in between. (b) $EC_{50}$ measurements of the toehold variants shown in (a). (c) Fold change of the toehold variants shown in (a). ON-state and leak for the (d) A-U toehold variants and (e) U-A toehold variants shown in (a). (f) Schematic detailing how 1-6 bp toeholds were introduced to a loop 7A background to test the importance of stacking in suppressing dynamic range. (g) $EC_{50}$ measurements of the toehold variants shown in (f). (h) Fold change of the toehold variants shown in (f). (i) $\Delta EC_{50}$ measurements of the toehold variants shown in (f) relative to polyA loops of the same overall spacer length (e.g. a 2 bp toehold introduces 4 additional nucleotides to the spacer region containing a 7A loop, so the $EC_{50}$ of an 11A polyA loop was used to calculate $\Delta EC_{50}$ for this particular comparison). Standard error propagation was performed by calculating the square root of the sum of the squares of the standard errors of the two comparisons. (j) ΔFold change of the toehold variants shown in (f) relative to the fold change of riboswitches with a polyA loop of the same overall length. (k) ON-state and leak for the A-U toehold variants. (l) ON-state and leak for the U-A toehold variants. Data in panels (b-e) and (g-l) are determined as described in **Methods**.

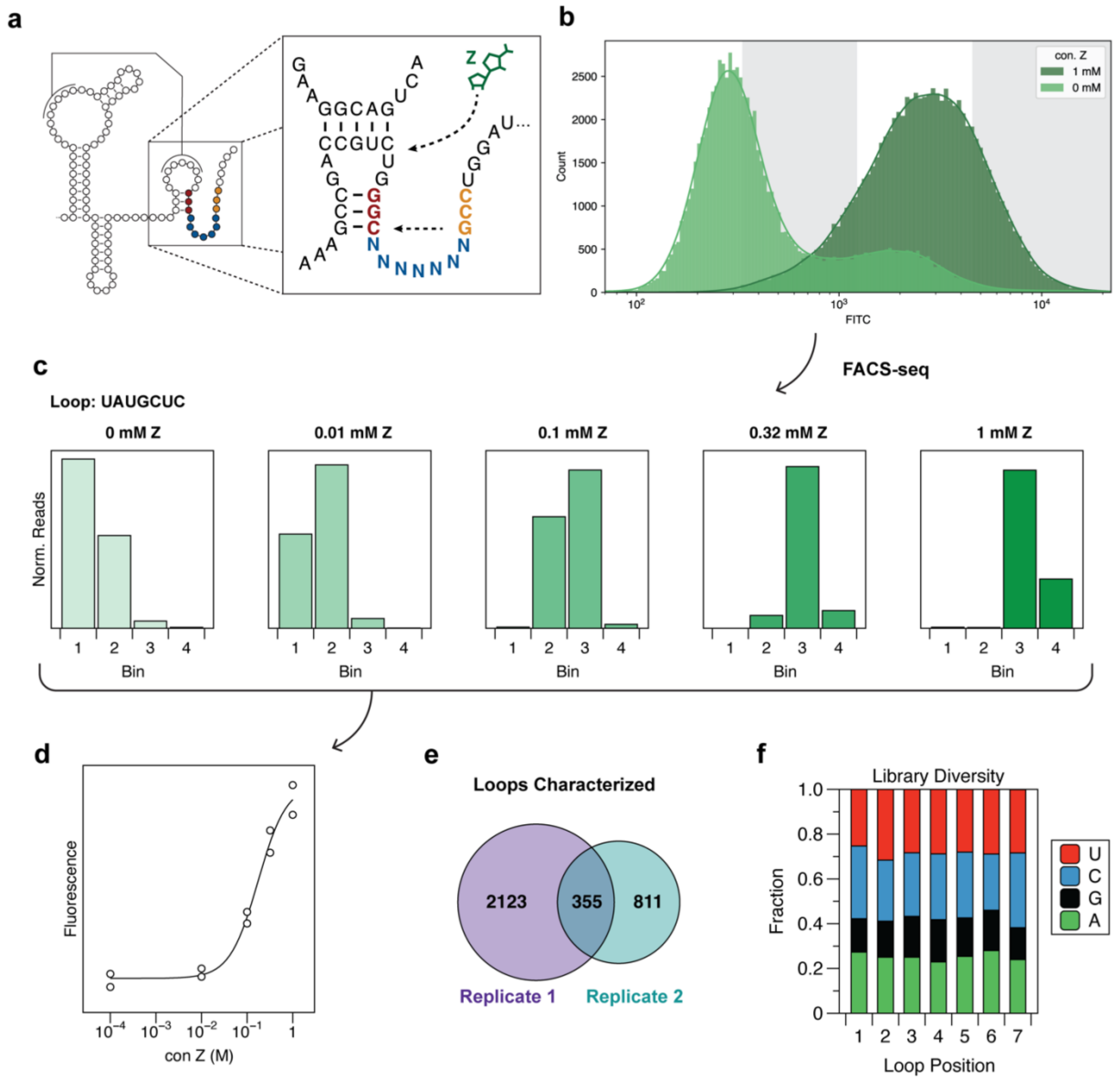


**Figure S9: FACS-Seq enables high-throughput characterization of riboswitch function.** (a) Schematic of random loop library design. (b) Random loop library fluorescence phenotype in the presence and absence of Z. White and gray alternating background indicate the width of bins used to sort the library by FACS. Bin positions were defined as described in Supplementary Data File 1. (c) An example loop sequence, UAUGCUC, demonstrating how low-resolution histograms resulting from the normalized reads per bin were obtained via FACS-seq. (d) Weighted geometric averages of the histograms in (c) were used to fit dose-response curves to obtain fold change and $EC_{50}$ measurements. (e) Two FACS-seq replicates were performed, each obtaining 2478 and 1166 unique loop sequences respectively with corresponding data that met the quality standards described in **Methods**. 355 loop sequences were characterized by both experiments. (f) Base identity fraction for each position within the random loop in the 1675 loops characterized by FACS-seq.

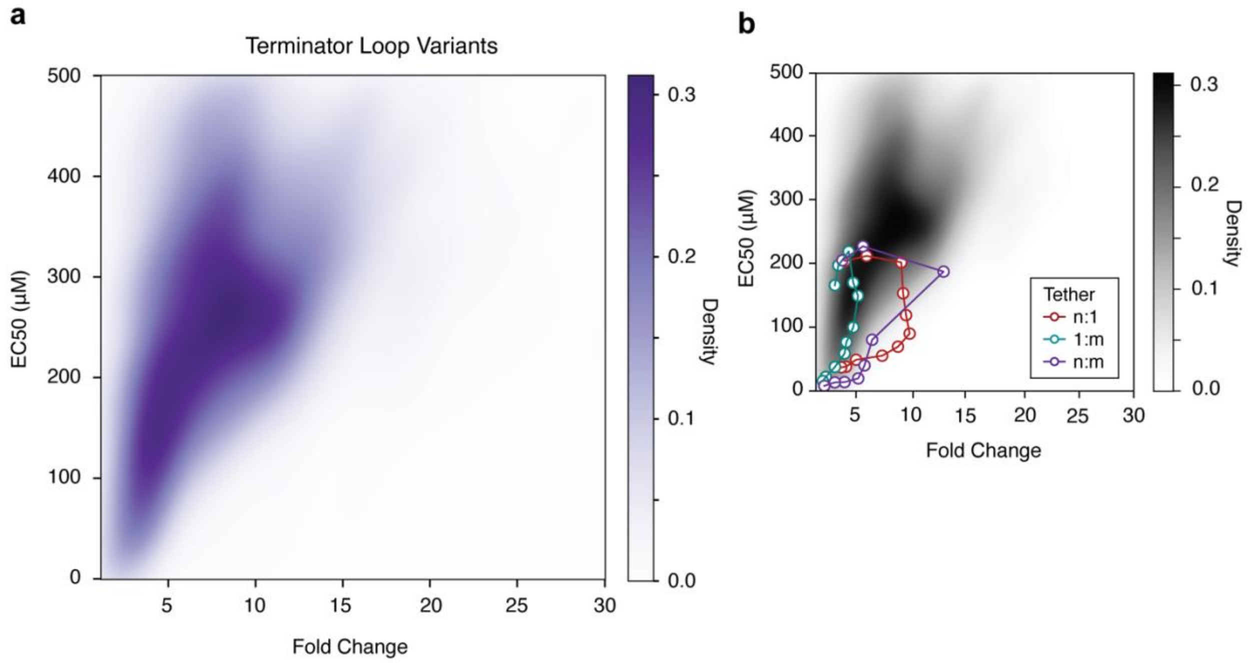


**Figure S10: High throughput FACS-Seq characterization of riboswitch fitness landscape mediated by terminator loop sequence.** (a) Density plot of the extracted $EC_{50}$ and fold change measurements obtained for all 3289 loop variants via FACS-seq. Density was calculated by performing a Gaussian kernel density estimate on extracted $EC_{50}$ and fold change values (see **Methods**). (b) Density plot from (a) overlaid with data from the curves observed for remote toeholds in Fig. 3.

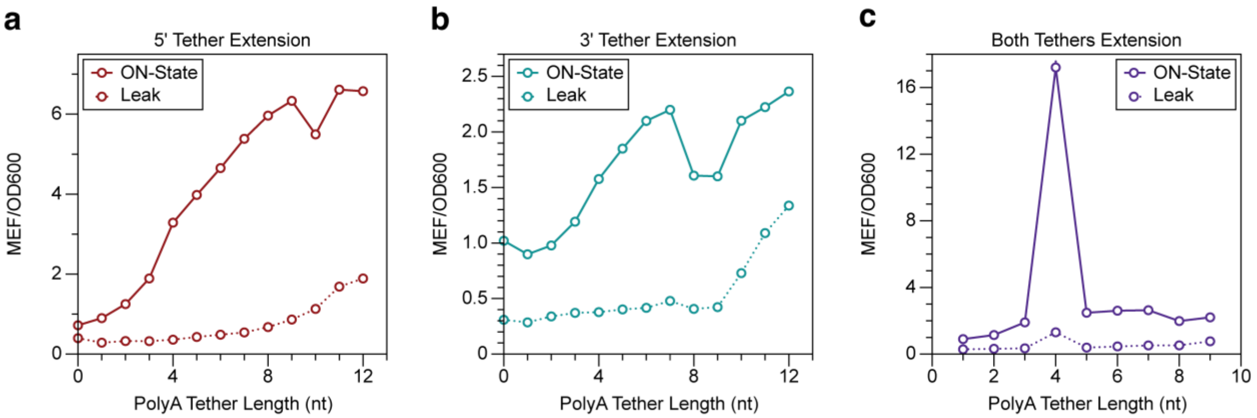


**Figure S11: ON-state and leak for remote toeholds from Figure 3.** ON-state ($F_{max}$) and leak ($F_{min}$) for (a) 5' leash extension remote toehold variants (n:1); (b) 3' leash extension remote toehold variants (1:m); and (c) both leash extension remote toehold variants (n:m). Bars for all panels indicate standard deviation. Data are determined as described in **Methods**.

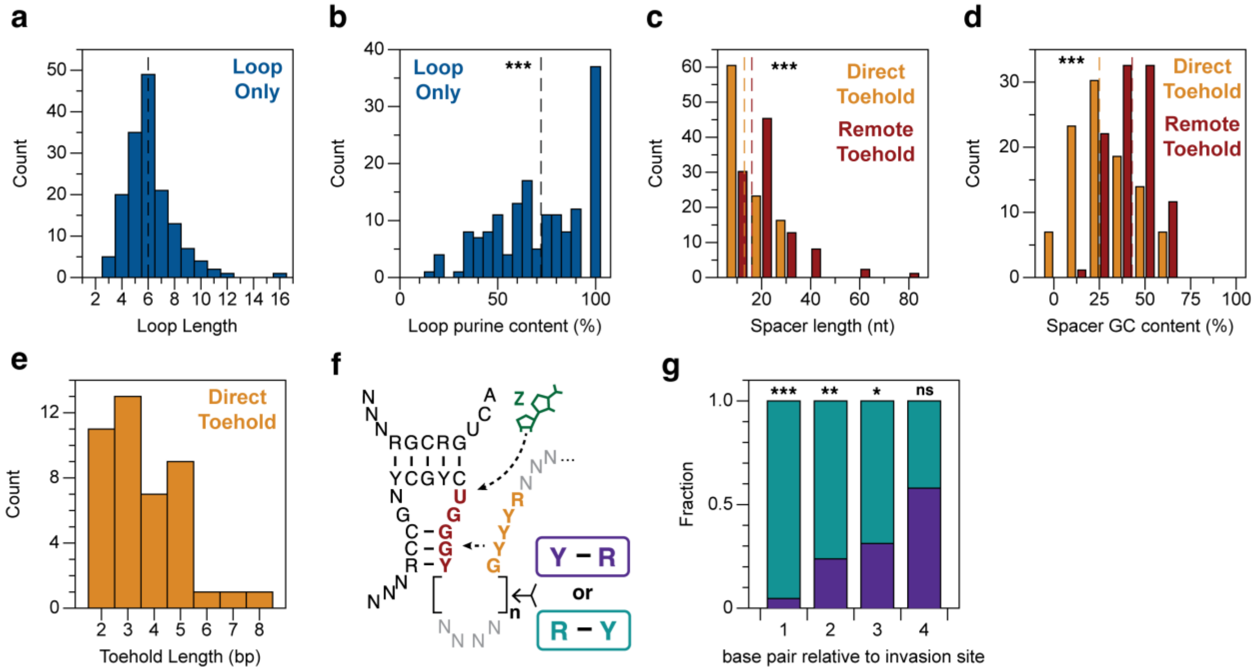


**Figure S12: Bioinformatic analysis of natural ZTP riboswitch EP architectures.** (a) Histogram of the length of loop-only EP loops. Dashed vertical line indicates median. (b) Histogram of the purine content (AG%) of loop-only EP loops. Dashed vertical line indicates median. *** indicates p-value <0.001 in a Wilcoxon Ranked-Sign test assuming a null hypothesis of 50%. (c) Histogram of the length of Direct Toehold (gold) and Remote Toehold (red) EP spacer regions. Dashed vertical lines indicates median value, and *** indicates p-value <0.001 in a Mann-Whitney U test demonstrating the two distributions are significantly different. (d) Histogram of the GC content of Direct Toehold (gold) and Remote Toehold (red) EP spacer regions. Dashed vertical lines indicates median value, and *** indicates p-value <0.001 in a Mann-Whitney U test demonstrating the two distributions are significantly different. (e) Toehold lengths observed in the 43 identified direct toehold EPs. (f) Schematic of the consensus structure of Rfam used to identify Direct Toeholds. (g) Analysis of the orientation of toehold base pairs from the 43 spacer regions with ≥2 bp direct toeholds. Stars indicate p-values <0.001 (***), <0.01 (**), <0.05 (*) in a chi-squared test assuming a 50% likelihood of either base pair orientation. For base pair positions 1 and 2 n=43, for position 3 n=32, and for position 4 n=19.

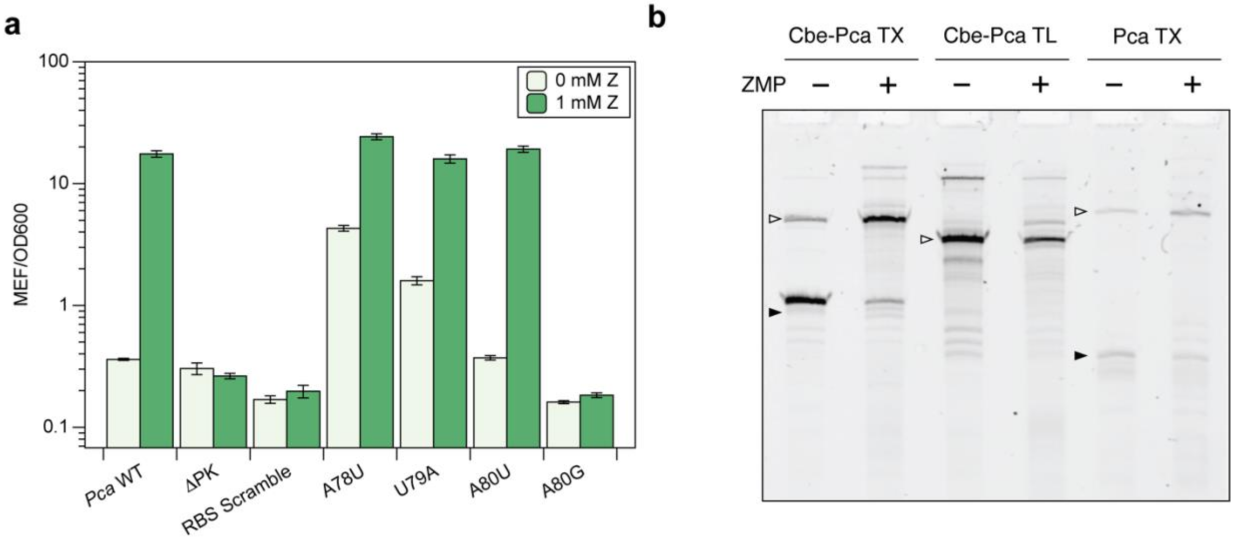


**Figure S13: Characterization of *Pca rhtB* and EP chimeras.** (a) Mutagenesis of *Pca rhtB* EP validates previously suggested switching mechanism (8). Deleting the 5' bases in the pseudoknot (ΔPK) completely breaks function, as does scrambling the RBS sequence. Disrupting base pairs in the anti-start codon (A78U, U79A) reveals that these base pairs are important for suppressing riboswitch leak. The mismatch between A80 and C103 is required for the riboswitch to turn ON. Mutating A80 to U, which maintains the mismatch, has no effect on function, while A80G, which allows a new base pair to form at this position, and converts the EP to an 8 bp direct toehold, breaks the riboswitch OFF to equivalent expression levels as the ligand non-binding ΔPK mutant. (b) In vitro transcription of chimeric *Pca* chimeric variants in the presence (1 mM) and absence of ZMP. *Pca TX* and *Cbe-Pca TX* show ligand-dependent increases in transcriptional readthrough, while no terminated band is visible for in either ligand condition for *Cbe-Pca TL*. Filled triangles indicate the terminated length, and open triangles indicate full-length product based on length of the linear template DNA. Source gel images of 2 replicates can be found in Fig. S14. Bars in panel (a) indicate average MEF/OD600 values over n = 9 replicates, with error bars indicating standard deviation.

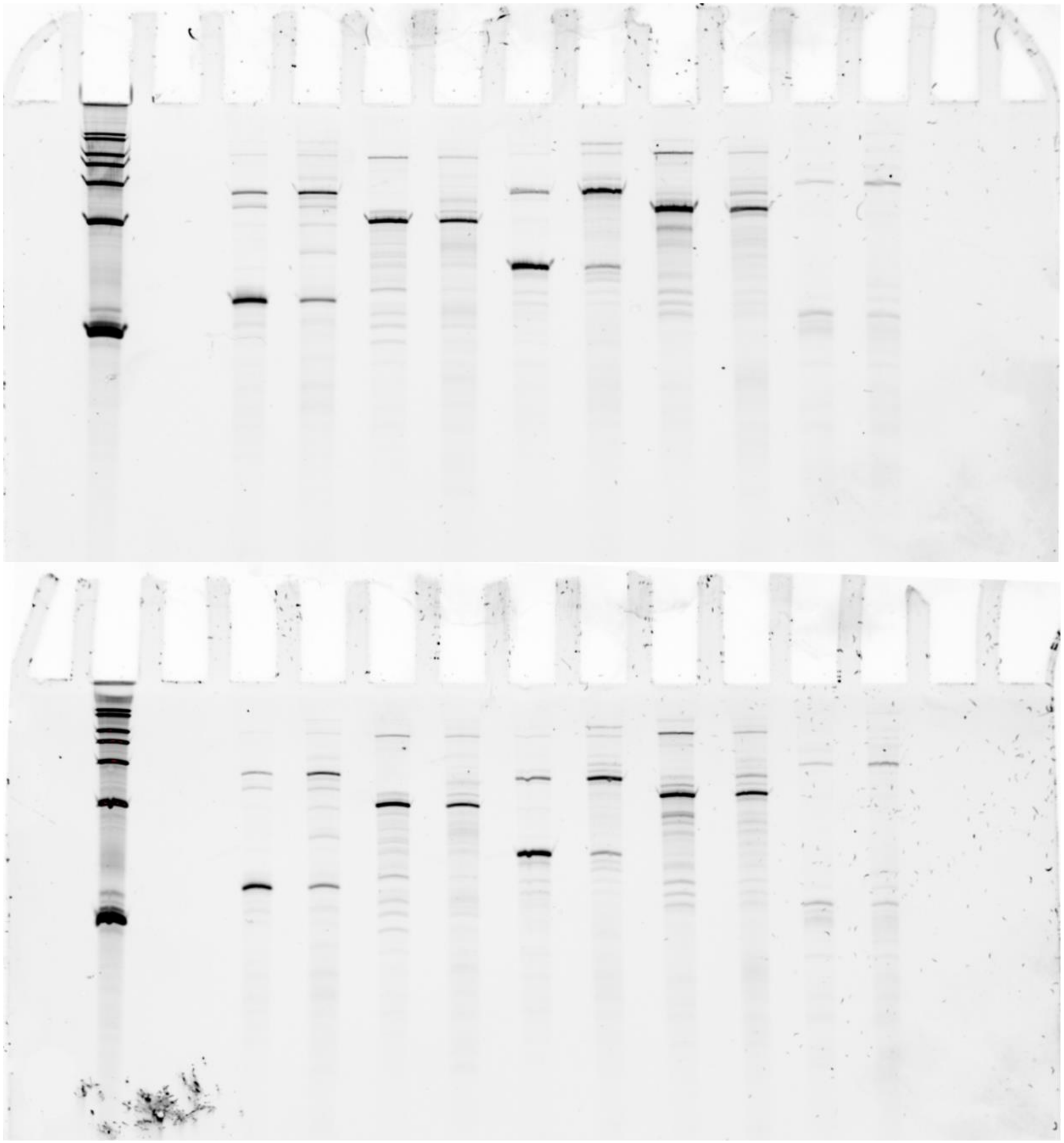

**Figure S14: Replicate source gels for Figures 5 and S13.** Samples from left to right are (1) Century™-Plus RNA Ladder (Invitrogen™), (2) Blank, (3) *Cbe pfl* (pJBL3907) 0 mM ZMP, (4) *Cbe pfl* (pJBL3907) 1 mM ZMP, (5) *Pca rhtB* (pJBL6931) 0 mM ZMP, (6) *Pca rhtB* (pJBL6931) 1 mM ZMP, (7) *Cbe-Pca* TX (pJBL7316) 0 mM ZMP, (8) *Cbe-Pca* TX (pJBL7316) 1 mM ZMP, (9) *Cbe-Pca* TL (pJBL7345) 0 mM ZMP, (10) *Cbe-Pca* TL (pJBL7345) 1 mM ZMP, (11) *Pca* TX (pJBL7346) 0 mM ZMP, (12) *Pca* TX (pJBL7346) 1 mM ZMP.

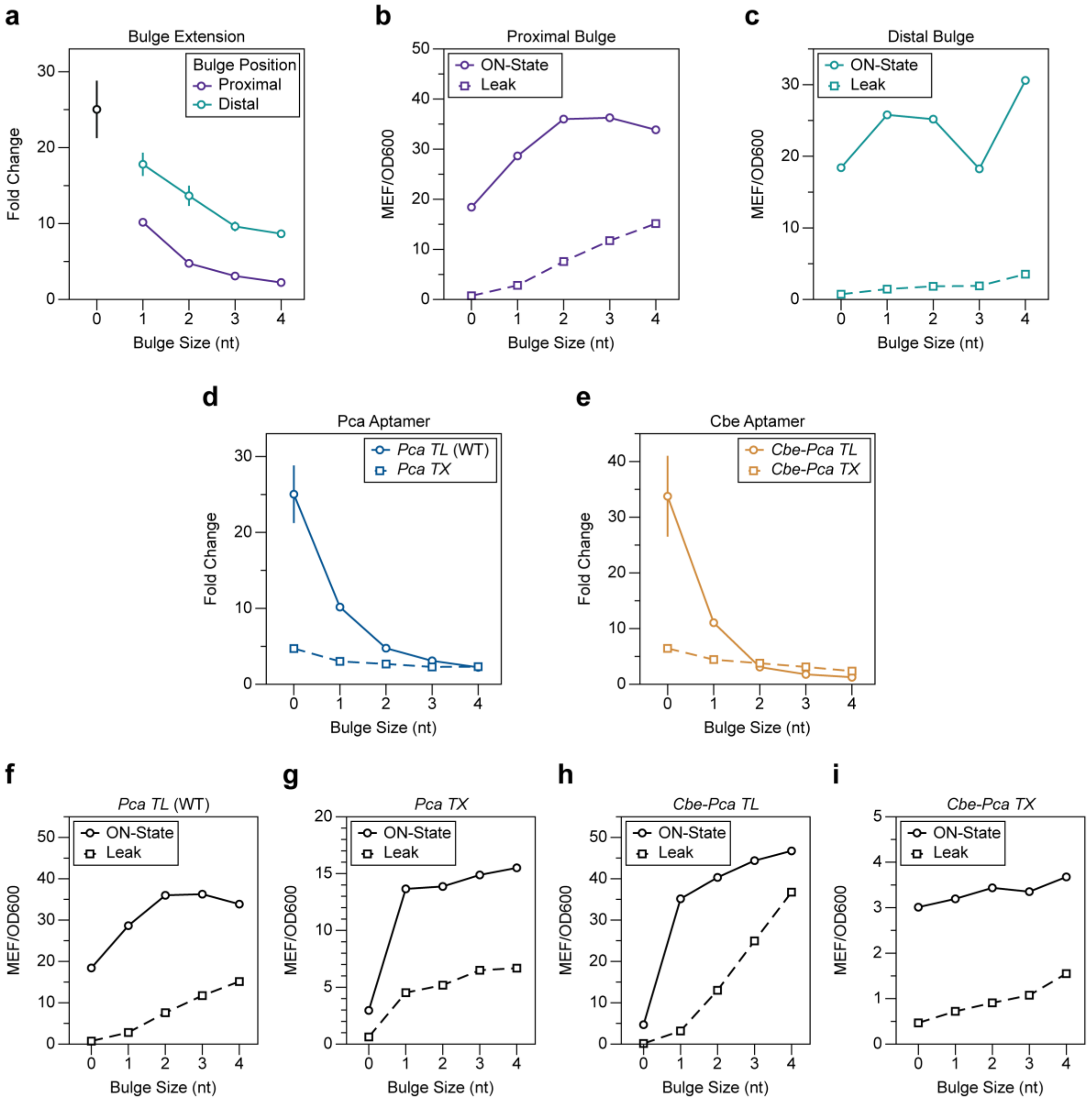


**Figure S15: ON-state and leak for bulge extension mutants in Figure 5.** (a) Fold change of *Pca rhtB* bulge mutants in Fig. 5C. (b) ON-state ($F_{max}$) and leak ($F_{min}$) for proximal bulge variants of *Pca rhtB*. (c) ON-state ($F_{max}$) and leak ($F_{min}$) for distal bulge variants of *Pca rhtB*. (d) Fold change of *Pca* AD bulge variants in Fig. 5E. (e) Fold change of *Cbe* AD bulge variants in Fig. 5F. (f-g) ON-state ($F_{max}$) and leak ($F_{min}$) for *Pca TL* (f), *Pca TX* (g), *Cbe-Pca TL* (h), and *Cbe-Pca TX* (i). Data are determined as described in **Methods**.

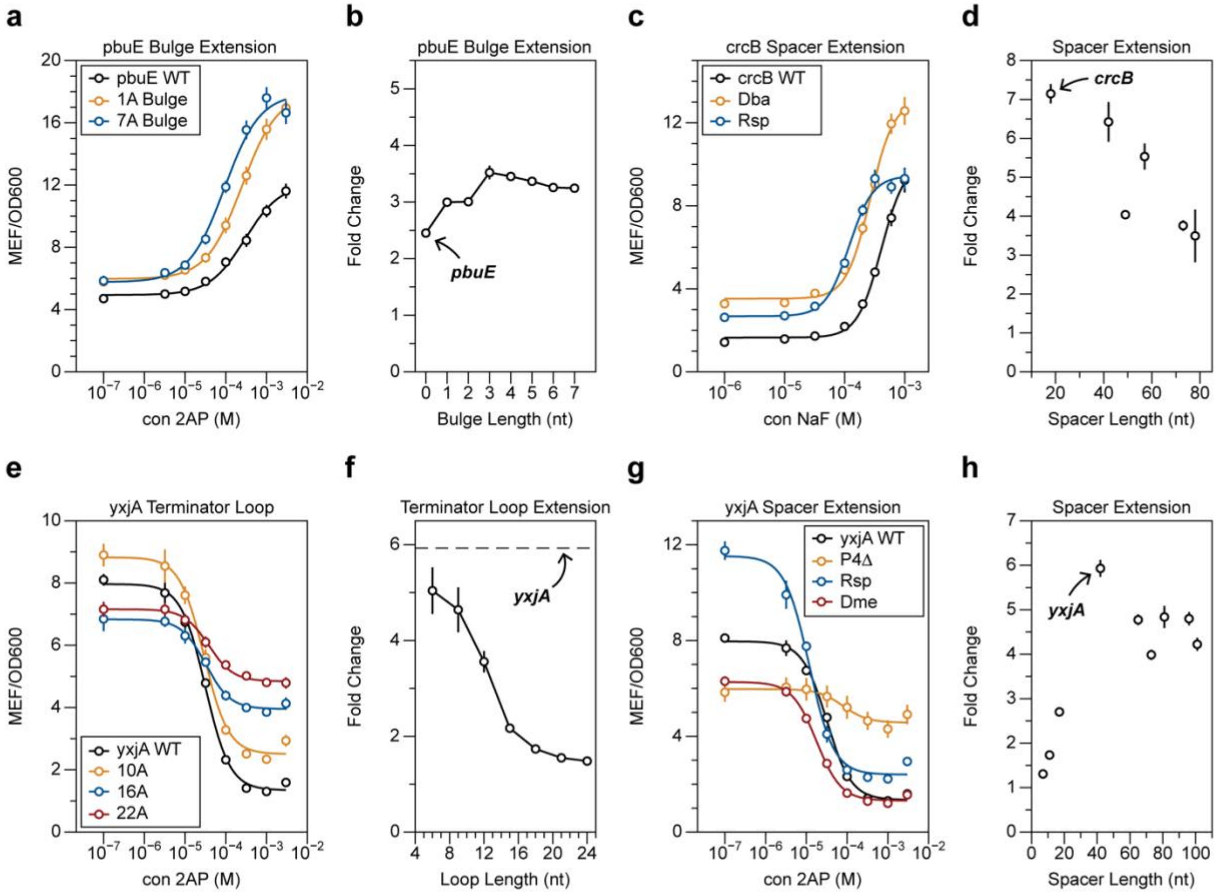


**Figure S16: Delaying EP nucleation enhances sensitivity for diverse transcriptional riboswitches.** (a) Select dose-response curves of *pbuE* bulge variants. (b) Fold change of *pbuE* bulge variants in Fig. 6B. Note that these fold change values are not directly comparable with those in (9), which were calculated by subtracting *E. coli* autofluorescence, leading to much higher calculated fold change values. (c) Select dose-response curves of *Bce* terminator extension variants. (d) Fold change of *Bce* terminator extension variants in Fig. 6D. (e) Select dose-response curves of *yxjA* terminator loop extension variants. (f) Fold change of *yxjA* terminator loop extension variants in Fig. 6F. (g) Select dose-response curves of *yxjA* P4 spacer extension/truncation variants. (h) Fold change of *yxjA* P4 spacer extension/truncation variants in Fig. 6G. Data in panels (a), (c), (e), and (g) represent dose response curves over n = 9 replicates, with error bars indicating standard deviation. Data are determined as described in **Methods**.

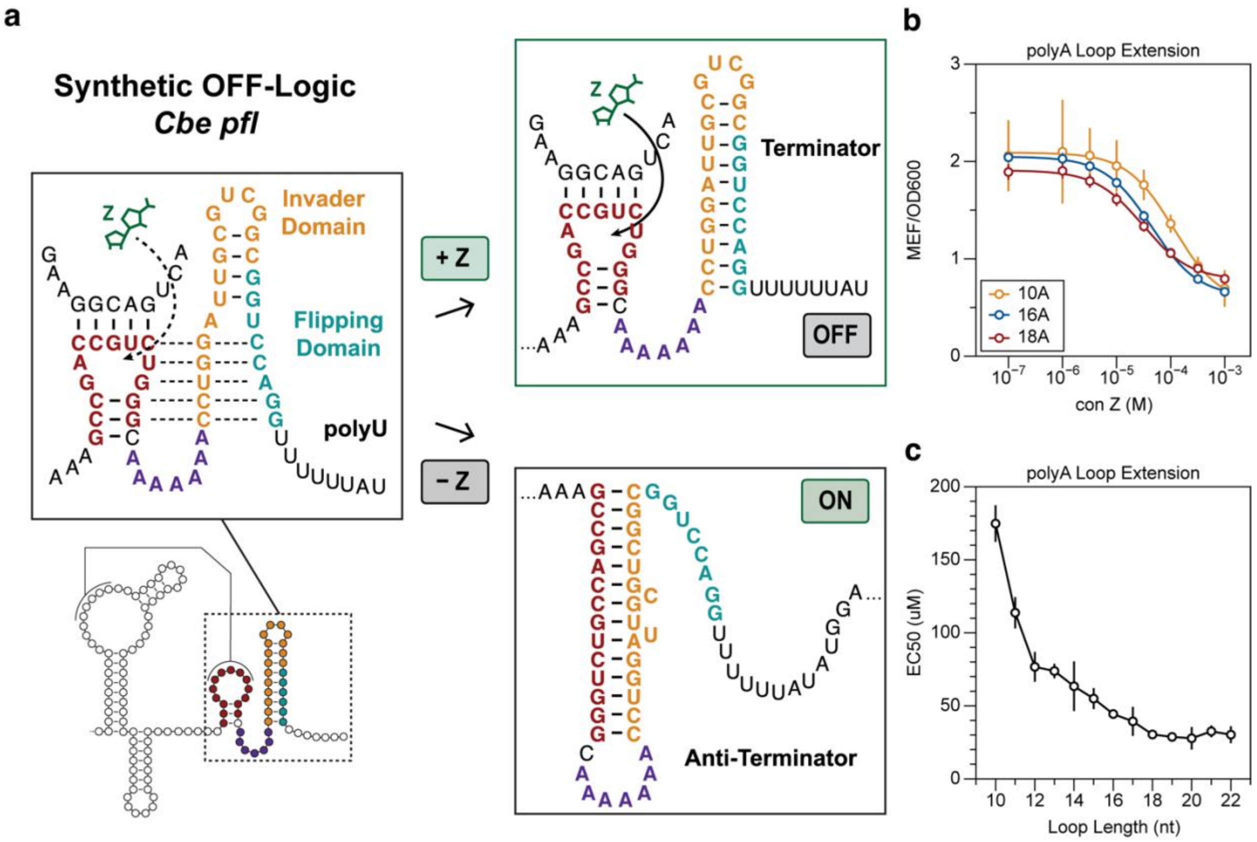


**Figure S17: Loop length tunes sensitivity in a synthetic OFF-logic ZTP riboswitch.** (a) Schematic detailing the synthetic OFF ZTP expression platform as originally described in (10). The presence of an additional synthetic flipping domain (teal) inverts the regulatory logic of the *Cbe pfl* riboswitch. (b) Select dose response curves of OFF-logic ZTP riboswitches with varying length polyA loops. (c) $EC_{50}$ measurements for varying polyA loop extension variants. Data in panel (b) represents dose response curves over n = 9 replicates, with error bars indicating standard deviation. Data in panel (c) are determined as described in **Methods**.

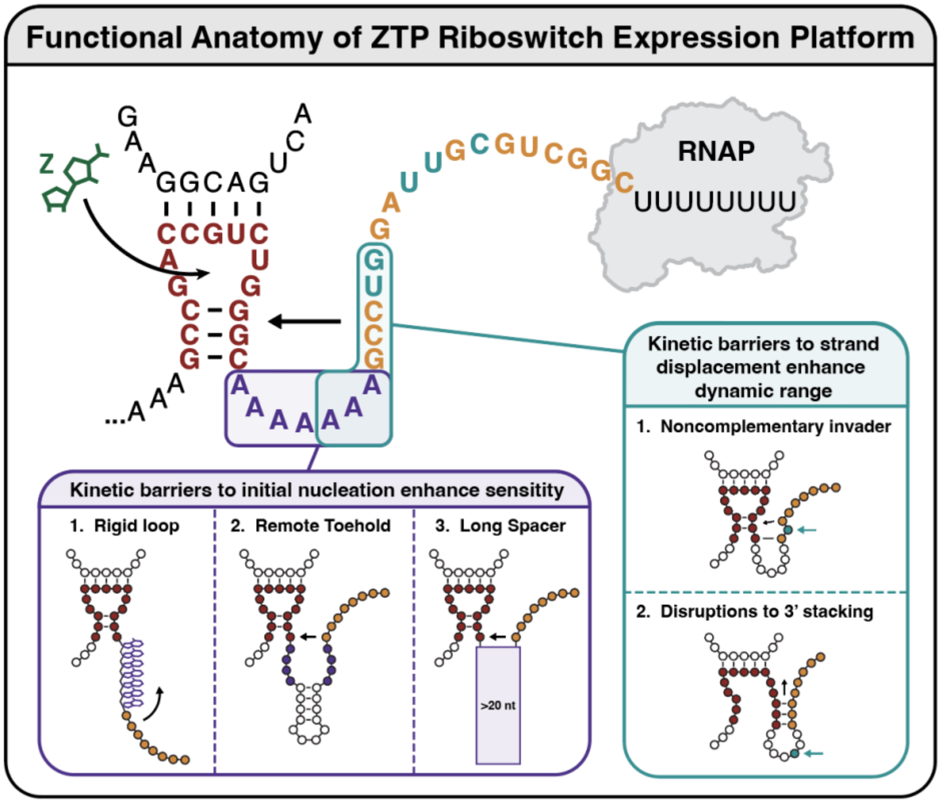


**Figure S18: Functional anatomy of the *Cbe pfl* ZTP riboswitch.** Combining the results of previous studies investigating how slowing EP strand displacement programs dynamic range (10) with the current study allows for a functional anatomy of the ZTP riboswitch expression platform. The *Cbe pfl* ZTP riboswitch EP features two overlapping functional domains. The first domain (purple) controls sensitivity by tuning the rate of initial strand displacement nucleation. This can be accomplished via A-A base stacking, remote toehold architecture, or simply by extending the length of the spacer region to consume time with transcription. The second functional domain (teal) controls dynamic range by tuning the rate of strand displacement. This can be accomplished by the presence of non-complementary bases (teal) in the invader region (gold), or by the disruption of stacking interactions within the 3' region of a terminator loop or toehold.

| Genome | GenBank Accession | Down-stream Gene | Full Sequence to polyU | Spacer Region Sequence | Spacer Region Class |
|---|---|---|---|---|---|
| Sharpea azabuensis strain DSM 20406 | FNYK010000 49.1 | IMPCH | AGGTAGTTATACGACTGGCGG AAGTGGAATTAACCACATGAA GTATAATCGTTGTAGAGCCGA CCGTCTGGGCAGGAGAAGCG CCAGGGGTAGTAGGCTCTTTT TTC | AGGAAGAA | Loop Only |
| Clostridium kluyveri | CP000673.1 | pfl | AAGGAATCGTATAACCGGCGG CAGTGGAATTAACCACAAGGA GTACGATTTTTAAAAGCCGAC CGCCTGGGCAATCAAAAAGTC CAGGTGGTCTTTTTGA | AATCAAAA A | Loop Only |
| Clostridium tyrobutyricum | CBXI0100000 31.1 | pfl | AATGAGTCGTATAACCGGCGG AAGTGGATATAACCACAGGGA GTACGATTTTTAAAAGCCGAC CGCCTGGGCAAACAAAAAGTC CAGACAGTCGGTCTTTTTGT | AAACAAAA A | Loop Only |
| Acidaminococcus intestini RyC-MR95 | CP003058.1 | ofa | AAAGCAGTGCAGGACTGACG GATAAGTGGAATTGACCACGT GCTCTGCATTGTGTAACTTGC CGACCGTCTGGGCAGAGAAAA GCAGCCCGGACGGTCCTTTTT CA | AGAGAAAA GCA | Loop Only |
| Desulfitobacterium metallireducens | CP007032.1 | pfl | AAACGGTTACATGACTGGCGG AAATATTGTAGACTTAACTACG ATGAAGTGTAACTGGAGAAAT GCCGACCGCCTGGGCGATTAT TTACAGGAATGTAAAATAATCG TTCAGGCGGTTTTTTAT | GATTATTTA CAGGAATG TAAAATAAT C | Direct Toehold |
| Deltaproteobacteria bacterium | MGSN010000 40.1 | idh | GACTTCTTCTGCAACTGACGG AATTAAGGTGGTTACCACCGG GGAGCTGGAGATATTTTATAA GCCGACCGTCTGGGCAGAAA GAGCTTGGCCAAAAGCTACGC AATTTCACAGCTCAGGCGGTT TTTTTA | AGAAAGAG CTTGGCCA AAAGCTAC GCAATTTC ACA | Remote Toehold |
| Lachnoclostridium phytofermentans | CP000885.1 | IMPCH | ATTACAATGTACGACTGGCGG AAACTGTAGGAGACTACAGGT GGGGATAACCCACAGGGAGT | CACCACCT TCGATGAT ATCCATGA | Remote Toehold |

| | | | | | |
|---|---|---|---|---|---|
| | | | ACATTTAATAACAATAAGAGCC GACCGCCTGGGCCACCACCTT CGATGATATCCATGATGGATA TCCGTTGGCAGTAGCTCAGGT GGTTTTTTT | TGGATATC CGTTGGCA GTA | |
| Roseburia sp. CAG:197 | HF999930.1 | IMPCH | ATTTAGTTATATGACTGACGGA ATTTATGTGGAAGAACCACGT GGAGTATAACATCAGGCAGTT TTTAACTGCTGGGAAAGGCAA TGTCTCTTCGGGACATCGCGG CAACGATTCATAGAAACGTTG CAATTGCCGACCGTCTGGGCA ACCTGCACATTTGAGGGCAAT GCAGTGCGCATTACCTTATAT GCTATTGTGTAAGTTCTCAGAT GTGTCGGCTTTTTTCA | AACCTGCA CATTTGAG GGCAATGC AGTGCGCA TTACCTTAT ATGCTATT GTGTAAGT T | Remote Toehold (lacks G108) |

**Table S1: Sequences of ZTP riboswitches used to generate chimeras in Fig. 4**

### SUPPLEMENTARY DATA FILES

Supplementary Data File 1: Excel file containing sequences of all riboswitch variants, all fluorescence data, MEF calibration, and FACS-seq information.

Supplementary code files along with descriptions can be found at https://github.com/LucksLab/Bushhouse_Riboswitch_Sensitivity_2024 .
